# Fibroblasts-derived extracellular vesicles contain SFRP1 and mediate pulmonary fibrosis

**DOI:** 10.1101/2022.12.22.521499

**Authors:** Olivier Burgy, Christoph H. Mayr, Beatriz Ballester Llobell, Arunima Sengupta, Déborah Schenesse, Christina Coughlan, Tanyalak Parimon, Peter Chen, Michael Lindner, Anne Hilgendorff, Matthias Mann, Ali Önder Yildirim, Oliver Eickelberg, Herbert B. Schiller, Mareike Lehmann, Gerald Burgstaller, Melanie Königshoff

## Abstract

Idiopathic pulmonary fibrosis (IPF) is a lethal and chronic lung disease characterized by aberrant intercellular communication, increased extracellular matrix (ECM) deposition, and destruction of functional lung tissue. Extracellular vesicles (EVs) accumulate within the lung in IPF, but their cargo and biological effects remain unclear. Here, we provide the entire the proteome of EV and non-EV fraction during pulmonary fibrosis, and functionally characterize their contribution to fibrosis. EVs were isolated by differential ultracentrifugation of bronchoalveolar lavage fluid (BALF) collected from mice challenged with bleomycin (or PBS as control) or culture supernatants from primary mouse lung fibroblasts. EVs were characterized by nanoparticle tracking analysis, Western Blotting, and quantitative mass spectrometry to define their proteome. EVs accumulation peaked at 14 days post-bleomycin instillation and correlated with decreased lung function. Label-free proteomics identified 107 proteins specific to fibrotic BALF-EVs. This signature was associated with wound healing, extracellular matrix organization, and cell motility. BALF-EVs from fibrotic lungs promoted fibrogenesis, including induction of ECM proteins in precision cut lung slices *ex vivo* and impaired alveolar epithelial cell stem cell function. Deconvolution using single cell RNA sequencing datasets revealed that fibroblasts are the major cellular source of BALF-EVs. EVs from fibroblasts were significantly enriched in Secreted Frizzled Related Protein 1 (SFRP1). In the lungs of patients with IPF, SFRP1 was significantly increased in mesenchymal cells. *Sfrp1* deficiency reduced the ability of fibroblast-derived EVs to potentiate bleomycin-induced lung fibrosis *in vivo* and led to a reduction in fibrosis marker gene expression. In sum, EVs carry specific protein cargos, such as SFRP1, to contribute to organ remodeling during fibrosis. Our data identified EVs transporting SFRP1 as a potential therapeutic target for IPF.

## Introduction

Fibroproliferative diseases can affect all tissues and organ systems. They represent a major health problem and are responsible for 45 % of deaths around the world (Wynn, 2007). Among them, idiopathic pulmonary fibrosis (IPF) is a chronic progressive and fatal fibrotic disorder of the lung (Martinez et al., 2017). Current therapies are limited to two approved drugs, pirfenidone and nintedanib, which slow down the progression of the disease but are unable to stop or reverse it (King et al., 2014; Richeldi et al., 2016). Thus, there is a major unmet clinical need for targeted therapies. Our understanding of the molecular mechanisms driving IPF initiation and progression is still limited, however, based on recent data, IPF is thought to be driven by repetitive insults to the lung epithelium that results in a local pro-fibrotic milieu within the lung where fibroblasts, the key effector cells in fibrosis, are activated and lead to increased alerted extracellular matrix (ECM) deposition (Martinez et al., 2017). The (re)activation of developmental signaling pathways, such as TGF-β or WNT leads to impaired cell to cell communication resulting in tissue fibrosis and scarring (Burgy et al., 2020; Burgy and Konigshoff, 2018; Fernandez and Eickelberg, 2012). The mechanisms involved in cellular crosstalk contributing to fibrosis, however, are still to be understood. In addition, whether their inhibition can mitigate fibrosis development and progression remains largely unknown.

Extracellular vesicles (EVs) have emerged as potent contributors to cellular crosstalk. They represent a group of membranous structures with a size range from 30 to 1000 nm depending on origin and are secreted by all cells (van Niel et al., 2018). They are classified into two major groups: exosomes which originate from intraluminal vesicles within multivesicular bodies, and ectosomes produced by budding from the plasma membrane (van Niel et al., 2022). EVs contain a wide array of cargo, from proteins to nucleic acids and lipids, and thus are major players in cellular crosstalk (Zomer et al., 2015). The diverse cargo transported by the vesicles mediate the biological activity of EVs, however their composition and distinct effects in fibrosis in general, and in pulmonary fibrosis in particular, is poorly understood (Burgy et al., 2020). We and others have recently demonstrated that EVs are increased in experimental lung fibrosis as well as in human PF (Martin-Medina et al., 2018; Njock et al., 2019). During fibrosis, EVs carry specific nucleic acids such as miRNAs, which can alter pro-fibrotic signaling (Njock et al., 2019; Parimon et al., 2019). Moreover, we found that EVs carry WNT5A, which potentiates profibrotic fibroblast function (Martin-Medina et al., 2018). Together, these data strongly support the notion that EV carry distinct cargo as impactful mediators of fibrosis, which might be amenable for therapeutic targeting.

To gain a deeper understanding of the altered EV cargo in pulmonary fibrosis, and potential novel therapeutic targets and biomarkers, we performed an unbiased analysis of the EV proteome in lung fibrosis initiation and progression. We found a distinct profile of proteins enriched specifically in EVs following bleomycin exposure and showed that fibrotic EVs impair lung epithelial stem cell function and modulate ECM deposition. We discovered that fibroblasts are key cells secreting EVs during lung fibrosis. These cells express high levels of Secreted Frizzled Related Protein 1 (SFRP1), a WNT family member protein that is one of the major proteins secreted by fibroblasts and enriched in fibrotic EVs. We demonstrate that EVs lacking SFRP1 have diminished pro-fibrotic properties *in vivo*, thus proving first evidence for novel therapeutic strategies targeting EV-linked proteins in pulmonary fibrosis.

## Materials and methods

### Human tissue and ethics statement

Primary human fibroblasts and pulmonary tissue from patients with IPF and patients without diagnosed chronic lung disease were obtained from the CPC-M bioArchive at the Comprehensive Pneumology Center (CPC Munich, Germany). The study was approved by the local ethics committee of the Ludwig-Maximilians University of Munich, Germany (Ethic vote #333-10). Written informed consent was obtained for all study participants.

### Ethics and study approval

The procedures involving animals in this study have been approved by the institutional animal care and use committee of the University of Colorado Denver (Aurora, CO, USA), the ethics committee of the Helmholtz Zentrum München and the Regierung von Oberbayern (Munich, Germany) and the “Comité d’Ethique de l’Expérimentation Animale du grand campus” of the University of Burgundy (Dijon, France) under the project references 115517(04)1E, AZ: 55.2-1-54-2532-88-2012 and APAFIS #26877. 8-week-old C57Bl/6J male mice were purchased from the Jackson Laboratory (Bar Harbor, ME, USA) or Charles River (Saint-Germain-sur-l’Arbresle, France). *S*ecreted frizzled-related protein 1 (*Sfrp1*) deficient mice SFRP1-2AKI (ref MGI:3796251) (Satoh et al., 2006) were ordered from MRC Harwell, Mary Lyon Centre, Harwell Campus, Oxfordshire. Here, the SFRP1 is a targeted (null) knock-out by deletion of exon1 by replacing it with a nuclear-localized lacZ KI cassette (IRES-T-LacZ-bpA) and a PGKneobpA (1700 bp) lox. The lacZ-neo cassette replaces exon 1 of the *Sfrp1* gene, abolishing gene function. The SFRP1-2AKI were received as frozen embryos (Stock ID: FESA:1734 Sales order reference: RQ217) and embryo transfer (FET) was performed in C57Bl/6J. The SFRP2 deletion was finally backcrossed, so that all progeny was SFRP2 homozygous wild type (+/+) for SFRP2. All the animals were housed in pathogen-free conditions, with environmental enrichment, in rooms maintained at constant temperature and humidity under a 12-h light cycle, and had access to food and water *ad libitum*.

### Bleomycin model

Mice were anesthetized with isoflurane and underwent orotracheal aspiration of bleomycin (Willow Birch Pharma, Taylor, MS, USA) with a single dose of 2.5 U/Kg. Control mice were injected with sterile 1X PBS. At the indicated time points, mice were anesthetized with a mixture of ketamine/xylazine before lung function testing (FX2-Flexivent®, SCIREQ, Montreal, QC, Canada). Three readings were taken for each animal. After exsanguination via the *vena cava*, broncho-alveolar lavage fluid (BALF) was collected, centrifuged (600g, 15min, +4°C) and cell-free BALF were stored at −80°C before being processed. Lung tissue were then harvested and either fixed with formalin for histology or flushed with 1X PBS and snap frozen in liquid nitrogen for RNA or protein extraction.

### Generation of murine PCLS

Precision cut lung slices (PCLS) were generated from healthy C57Bl/6J mice as previously described (Uhl et al., 2015). Briefly, using a syringe pump, the mouse lungs were filled via a tracheal cannula with 1.5% (w/v) warm, low gelling temperature melting point agarose (Sigma Aldrich, Saint-Quentin-Fallavier, France) in sterile DMEM medium (Dutscher, Bernolsheim, France), supplemented with antibiotics (Dutscher). Afterwards, the lungs were removed and transferred on ice in cultivation medium for 10 min to allow the agarose to solidify. Each lung lobe was separated and cut with a VT1200S vibratome (Leica, Nanterre, France) in 300 μm thick to generate 4mm diameter punches. PCLS were cultivated in DMEM supplemented with 0.1% FBS (Dutscher) and antibiotics. 24h after slicing, the lung tissue was exposed to EVs (BALF-EVs from mice with bleomycin-induced lung fibrosis or control mice. 1:1000 cell to vesicle ratio) for seven additional days. Media with EVs was replenished every 72h. Tissue viability was monitored by WST-1 assay (Sigma) as described (Uhl et al., 2015).

### Primary cells

Lung epithelial cells were isolated as previously described (Mutze et al., 2015; Ng-Blichfeldt et al., 2019) with slight modifications. Mice were anesthetized (ketamine/xylazine) before exsanguination by cutting the *vena cava*, and the lungs were flushed with PBS. Mouse lungs were intratracheally inflated with dispase (BD Bioscience) followed by 300-μl instillation of 1% low gelling temperature agarose (A9414, Sigma-Aldrich). Lungs were excised, minced and filtered through 100-, 20-and 10-μm nylon meshes (Sefar). White blood cells were depleted with CD45, and epithelial cells were selected using CD326 (EpCAM) magnetic beads (respectively 130-052-301 and 130-105-958, Miltenyi Biotec), according to the manufacturer’s instructions. EpCAM+ cells were resuspended in DMEM containing 2-mM l-alanyl–L-glutamine dipeptide (Gibco), 100-U·ml-1 penicillin and streptomycin (Sigma-Aldrich), 3.6-mg·ml-1 glucose (AppliChem) and 10-mM HEPES (Life Technologies) until further use.

Primary mouse lung fibroblast (pmLF) isolation was performed as previously described (Mummler et al., 2018). Sfrp1 deficient or control mice were anesthetized (ketamine/xylazine) before exsanguination by cutting the vena cava. After flushing, lung tissue was harvested and dissected lung lobes were placed in ice cold sterile PBS in a 6-well plate. Using sterile scalpels, the lobes were dissected into small 1-2mm pieces and immediately transferred into a 50ml falcon containing collagenase 1 (diluted 5mg per 50μl of 1X PBS). Collagenase digestion was carried on at 37°C at 400 rpm for 1 hour and next transferred into a 70μm-filter placed on a fresh falcon tube. Using the head of a syringe pistol the digested lung pieces were scratched onto the filter and rinsed thoroughly with sterile 1X PBS. The final suspension was centrifuged for 5 min at 400 rpm at 4°C. Lastly, the supernatant was discarded and the pellet was resuspended in fresh cell culture medium (20% FBS and 100U/ml of penicillin/streptomycin supplemented DMEM-F12 media) and cultured under standard cell culture conditions. To produce culture media for EV concentration, pmLF were cultivated in 175 cm^2^ flasks in DMEM supplemented with EV-free FBS (System Biosciences, Palo Alto, CA, USA). Importantly, cells were used at passages less than 12 and did not reach more than 90% confluency and viability (Trypan blue) was above 95%. Conditioned media from 48h of culture were collected from three independent clones of wild-type or *Sfrp1^-/-^* pmLF.

### Organoid assay

Organoids were cultured as previously described (Hu et al., 2020; Lehmann et al., 2020; Ng-Blichfeldt et al., 2019) with some modifications. MLg mouse lung fibroblasts (CCL-206, ATCC) were proliferation-inactivated by incubation in culture medium containing 10-μg·ml-1 mitomycin C (Merck) for 2 h, followed by three washes in warm PBS (Life Technologies), and let to recover in culture media for at least 1 h; 10,000 EpCAM+ cells suspended in 50-μl growth factor-reduced Matrigel (Corning) were diluted 1:1 with 10,000 MLg cells in 50-μl DMEM/F12 containing 10% (w/v) FBS and seeded into 96-well imaging plates (Corning). Cultures were maintained in DMEM/F12 containing 5% FBS, 100-U·ml-1 penicillin/streptomycin, 2-mM L-alanyl-l-glutamine dipeptide (Gibco), 2.5-μg·ml-1 amphotericin B (Gibco), 1× insulin-transferrin-selenium (Gibco), 0.025-μg·ml-1 recombinant human EGF (Sigma), 0.1-μg·ml-1 cholera toxin (Sigma-Aldrich), 30-μg·ml-1 bovine pituitary extract (Sigma-Aldrich) and 0.01 μM of freshly added all-trans retinoic acid (Sigma-Aldrich) on top of the matrigel; 10-μM Y-27632 (Tocris), a Rho-associated kinase (ROCK) inhibitor, was added to the media for the first 48 h of culture. EVs (1: 1000 epithelial over vesicle ratio) were added to the organoid media from Day 0 and in every culture media exchange. Media were refreshed every other day. Microscopy for organoid quantification at Day 14 was performed using a microscope Axiovert 40 (Zeiss). Individual organoids with a minimum size of 50 μm were manually counted by two investigators in a blinded manner. Organoid forming efficiency was calculated by dividing the number of organoids formed at D14 by the number of epithelial cells plated for the assay.

### EV concentration

EVs were concentrated from cell-free BALF and culture media under sterile conditions by differential ultracentrifugation using an already described protocol (Coughlan et al., 2020; Martin-Medina et al., 2018). In brief, samples were first centrifuged at 600g, 10min, 4°C to remove cells and cellular debris. Then, the supernatant was subjected to a first centrifugation at 10 000g, 30min, 4°C to pellet large vesicles and apoptotic bodies. The corresponding supernatant was centrifuged at 110 000g for 2h30 at 4°C to pellet so-called EVs. At this step, the supernatant (termed here EV-free SN) was saved and concentrated by Amicon Ultra-0.5 centrifugal filters (Merck Millipore). The pelleted EVs were washed in cold, 0.1 μm-filtered 1X PBS and centrifuged 110 000g, 2h30, 4°C. Finally, EVs were re-suspended in 200μl of 0.1μm-filtered 1X PBS. In this study, we have used two separate ultracentrifugation setting: Thermo Scientific Sorval WX Ultra 90 ultracentrifuge with T-647.5 and TFT-80.2 fixed-angle rotors (for characterization and proteomics) or Beckman Coulter L-80 ultracentrifuge with Type 45 and Type 50.4 2 fixed-angle rotors (for functional testing). Before sending out for label-free proteomic, EVs underwent size exclusion chromatography with a cutoff of 30nm, according to the manufacturer’s instruction (Exo-spin™, Cell Guidance Systems, Cambridge, UK).

### Nanoparticle Tracking Analysis

EV samples were characterized for size distribution and particle concentration by nanoparticle tracking analysis using a NanoSight NS300 system equipped with a green laser and a software version: NTA 3.2 Dev Build 3.2.16 (Malvern Panalytical, Malvern, UK). Frozen EVs were gently thawed and diluted in a final volume of 500μl of 0.1μm-filtered 1X PBS. Diluted samples were injected using a 1ml syringe at constant flow (25μl/min). Absence of air bubbles in the analysis chamber was confirmed before each measurement. Vesicles were tracked by a high frame rate camera (sCMOS camera, 25 frames per second) over a 30 sec time lap for a total of 5 replicates. Image acquisition and analysis were performed for all samples at constant temperature (22±0.1°C) with a set viscosity (1.05 cP) as well as fixed camera level (14) and detection threshold (6). Values from the five readings were manually checked to identify potential outliers.

### Proteomic analysis

Separated EV were dissolved in Triton X-100 based lysis buffer (Hepes 50 mM pH 7.4, NaCl 150mM, EDTA 5 mM, Triton X-100 0,5 %) and protein amount was measured using a Lowry based method (DC Protein Assay Kit 5000111, Bio-Rad). 10μg protein per sample were subjected for protein digestion and peptide purification using the iST label free sample preparation kit (PreOmics GmbH). From the EV-free supernatant, proteins were precipitated overnight at −20°C after mixing the supernatant with ice-cold acetone (1:4). Precipitated proteins were pelleted at 4000g at 4°C for 10min, the supernatant discarded and the protein pelleted air-dried for 10min. Proteins were resuspended in 6M GmdCl buffer (6M guanidinium hydrochloride, 100mM Tris-HCl pH 8, 10mM TCEP, 50mM CAA), incubated at 95°C shaking for 10min, cooled down and sonicated for 10 cycles at the Bioruptor (30sec pulse + 30 sec pause). Protein concentration was measured and 20μq were diluted 1:10 with digestion buffer (10% ACN, 25mM Tris) and digested with Trypsin/LysC 1:50 (enzyme/protein) overnight at 37°C. Samples were acidified to 1% TFA and peptides enriched using SDS-RPS stage tips as previously described (Kulak, Mann 2014). Approximately 1 μg of peptides were separated in 1 h gradients with reverse-phase chromatography being performed with an EASY-nLC 1000 ultra-high pressure system (Thermo Fisher Scientific), which was coupled to a Q Exactive Mass Spectrometer (Thermo Scientific) as previously described (Mayr et al., 2021).

### Processing of the scRNAseq dataset

Mass spectrometry (MS) raw files were processed using the MaxQuant software (Cox and Mann, 2008). As previously described (Schiller et al., 2015), peak lists were searched against the human UniProt FASTA database (November 2018), and a common contaminants database (247 entries) by the Andromeda search engine (Cox et al., 2011). Statistical and bioinformatics operations (such as normalization, imputation of missing values, annotation enrichment analysis, correlation analysis, hierarchical clustering, principal component analysis, and multiple-hypothesis testing corrections) were run with the Perseus software package (version 1.6.2.3.) (Tyanova et al., 2016). To determine compartment and cell type specific expression of EV specific proteins found by the MS analysis, the proteins were compared with a publicly available single cell RNA-sequencing dataset of whole lung from PBS and bleomycin treated mice sampled with Dropseq (Strunz et al., 2020). Data was analyzed using scanpy, a python package for the exploration of single-cell RNA-seq data (Wolf et al., 2018). Different time points of bleomycin treatment were pooled into one category to compare PBS and bleomycin specific expression of POIs. To determine cell type specific expression of POIs in the Venn diagram, scanpy’s tl.rank_genes_groups function was used to calculate significant cell type specific marker genes that were then matched with the EV specific POIs. Scoring of EV protein gene signatures on compartments and cell types was done using scanpy’s tl.score_genes function, which compares the average expression of signatures genes with randomly chosen reference genes. To determine the time component of Sfrp1 expression a recently published scRNAseq paper on mesenchyme enriched from bleomycin treated mouse lungs was used (Mayr et al., 2022).

### Western blot

Cells, lung homogenates and EV-Pellets were lysed in Triton X-100 lysis buffer supplemented with protease and phosphatase inhibitors (Roche Diagnostics, Mannheim, Germany) and protein concentrations were quantified using a modified Lowry assay (Bio-Rad). Samples were concentrated if necessary, by Amicon Ultra-0.5 centrifugal filters (Merck Millipore, Amsterdam, The Netherlands). Reducing conditions (4x Laemmli loading buffer: 150 mM Tris HCl, 275 mM SDS, 400 mM dithiothreitol, 3.5% (w/v) glycerol, 0.02% bromophenol blue) were used. For EV-Pellets or EV-free SN, constant amount of the fractions (1/5) were loaded into the gel, whereas constant protein amount (30 μg) was used for comparison of pmLFs or lung tissue. Samples were loaded into 10-12% SDS-PAGE gel and transferred to a nitrocellulose membrane that was blocked afterwards in PBS supplemented with 0.1% Tween20 (PBS-T) and 8% non-fat milk. Blocked membranes were incubated with primary antibodies directed against TGS101 (HPA006161, Sigma Aldrich), Nidogen-1 (ab14511, Abcam), SFRP-1 (ab126613, Abcam), α-SMA (ab5694, Abcam), AGER (ab3611, Abcam), QSOX (LS-C802615, Life Span), EGFR (4267, Cell Signalling), HSC70 (sc-7298, Santa Cruz Biotechnology), β-actin (A5316, Sigma) and GAPDH (sc-47724, Santa Cruz Biotechnology) at dilution of 1:1000 in PBS-T + 5% non-fat milk. Finally, membranes were incubated with Peroxidase-AffiniPure Goat anti-rabbit and anti-mouse IgG (Jackson ImmunoResearch, UK). The signal was detected by enhanced chemiluminescence reagents (Immobilon Crescendo Western HRP substrate, Merk) was imaged with a ChemiDoc MP Imaging System (Biorad). Ponceau S staining (Sigma Aldrich), HSC70, β-actin and GAPDH served as loading controls.

### Quantitative PCR

RNA were extracted from PCLS using an adapted version of the ZYMO quick-RNA protocol (Zymo Research, Orange, CA, USA). In brief, PCLS were incubated in 350ul TRIzol™ reagent (Thermo Fisher Scientific) for 30min on ice. After homogenization (Ultra-TURRAX), samples were centrifuged at 800g, 5min. Supernatant was mixed 1-1 (v/v) with 100% ethanol and loaded into a Zymo-Spin™ IIICG Column. The rest of the protocol was followed according to the manufacturer’s instructions. cDNA was prepared from 100ng of RNA using iScript™ cDNA Synthesis Kit (Bio-Rad) in a Veriti™ 96-Well Thermal Cycler (Applied Biosystems). Quantitative RT-PCR was performed using SYBR green Advanced Master Mix and run on a ViiA 7 Real-Time PCR System (all from Applied Biosystems). Primers were in-house designed (see table 1) and synthesized by Eurofins Genomics. Data were normalized to the expression of *Hprt* using the following calculation: CT *Hprt* – CT gene of interest.

### Histology

Formalin-fixed paraffin embedded (FFPE) sections were prepared and dewaxed by xylene and decreasing concentration of ethanol before staining with hematoxylin and eosin or picrosirius red (PSR). PSR-stained sections were observed under polarized light on an Axio Scope.A1 (Zeiss) and pictures captured with a Gryphax camera (Jenoptik). Briefly, ten random fields were imaged with constant parameters among the samples. High resolution images were then analyzed using ImageJ (Bethesda, MD, USA).

### Immunofluorescence staining and microscopy

Formalin-fixed paraffin-embedded (FFPE) lung tissue sections from PBS- and Bleomycin-treated mice, as well as from healthy donors and IPF patients, were first placed in an incubator at 60°C for an hour, which was followed by tissue deparaffinization process. Using a Microm HMS 740 Robot-Stainer (Thermo Fisher Scientific), the slides containing the tissue sections were automatically incubated with Xylene (2x, 5min.), 100% EtOH (2x, 2min.), 90% EtOH (1x, 1min.), 80% EtOH (1x, 1min.), 70% EtOH (1x, 1min.), and distilled water (1x, 30sec.). Next, the tissue section containing slides were placed in R-Universal buffer (Aptum Biologics) followed by transfer to an antigen retrieval buffer containing pressure chamber (2100 Retrieval, Aptum Biologics) After 30 mins inside the pressure chamber, the slides were washed once in 1X Tris buffer for 10 min and then incubated in 5% BSA in PBS for 40 mins at room temperature. Subsequently, tissue sections were stained with primary antibodies overnight at 4°C under humid conditions. The following primary antibodies were used: anti-SFRP1 (Abcam, #ab126613, 1:100) and alpha-Smooth Muscle Actin (SMA) (Sigma-Aldrich, A5228, 1:500). Cell nuclei were stained with DAPI (40,6-diamidino-2-phenylindole, 1:2,000; Sigma-Aldrich). Next day, the slides were washed twice in 1X PBS for 10 min, and further incubated with fluorescently-labeled secondary antibodies for 2 hours at room temperature under humid conditions. The following fluorescently labeled secondary antibodies were used: donkey anti-rabbit IgG Alexa Fluor-568 (1:500; Invitrogen) and donkey anti-mouse IgG Alexa Fluor-488 (1:500; Invitrogen). Following two additional washes, slides were then counterstained with DAPI for 1 hour at room temperature, washed again two times with 1X PBS for 10 min and subsequently dried at room temperature. Finally, tissue slides were mounted (Dako mounting medium) and kept in the dark at 4°C until further analysis. Images were acquired with an upright fluorescence microscope (AxioImager (Zeiss)), equipped with and Axiocam and using the following objectives: Plan-Apochromat 20×/0.8 M27 and Plan-Apochromat 63×/1.4 M27.The automated microscopy system was driven by ZEN2009 (Zeiss) software. The final images were cropped and adjusted for contrast and brightness by using the ZEN2012 (Zeiss) software.

### Statistical analysis

All data are expressed as mean ± SD and analyzed with GraphPad Prism 8 software (La Jolla, CA, USA). Normal distribution of the data distribution was determined by Kolmogorov-Smirnov testing with Lilliefors’ correction before applying a parametric test. Non-parametric Mann-Whitney test was used with comparison between groups when data did not follow a normal distribution. For comparison of more than 2 groups, one-way ANOVA was used followed by Dunnett’s post-hoc test. For correlation study, non-parametric Spearman test was used. For each comparison, two-tailed P value is indicated.

## Results

### EVs accumulate during active fibrosis and exacerbate its mechanisms

We performed an in-depth longitudinal time course of EV numbers and secretion during lung fibrosis initiation, progression and resolution (figure 1A). To this end, we applied a well-known experimental mouse model of lung fibrosis (Bonniaud et al., 2018; Jenkins et al., 2017). C57Bl/6J mice were challenged with a single orotracheal instillation of bleomycin (D0), which leads to an inflammatory phase (D3-7) followed by fibrosis (D10-21), and late resolution of the fibrotic lesions (D28-56) (Fig 1A). Pulmonary fibrosis was assessed by lung function and hydroxyproline content of the lung tissue, demonstrating active fibrosis at D14 (figure 1B, 1C), consistent with previous reports (Fernandez et al., 2016). Broncho-alveolar lavage fluid (BALF) was collected at the different time points post-bleomycin (from D3 to D56) and BALF-derived EVs (BALF-EVs) were concentrated by differential ultracentrifugation and quantified using nanoparticle tracking analysis (NTA). As shown in figure 1D, the amount of EVs was consistently increased in BALF samples at all-time points after bleomycin challenge compared with saline samples, with a peak at D14. Consistent with previous reports, these vesicles have a median size ranging from 103 to 190 nm in diameter (95% confidence interval) (figure 1E). Importantly, the quantity of EVs present at the BALF level correlated inversely with the lung function of mice exposed to bleomycin, further suggesting that the number of EVs is closely linked to profibrotic processes in the lung (figure 1F). Therefore, we focused our downstream analysis on BALF-EVs at D14 post-bleomycin exposure representing the phase of active fibrosis and the peak of EV accumulation.

**Figure 1.**
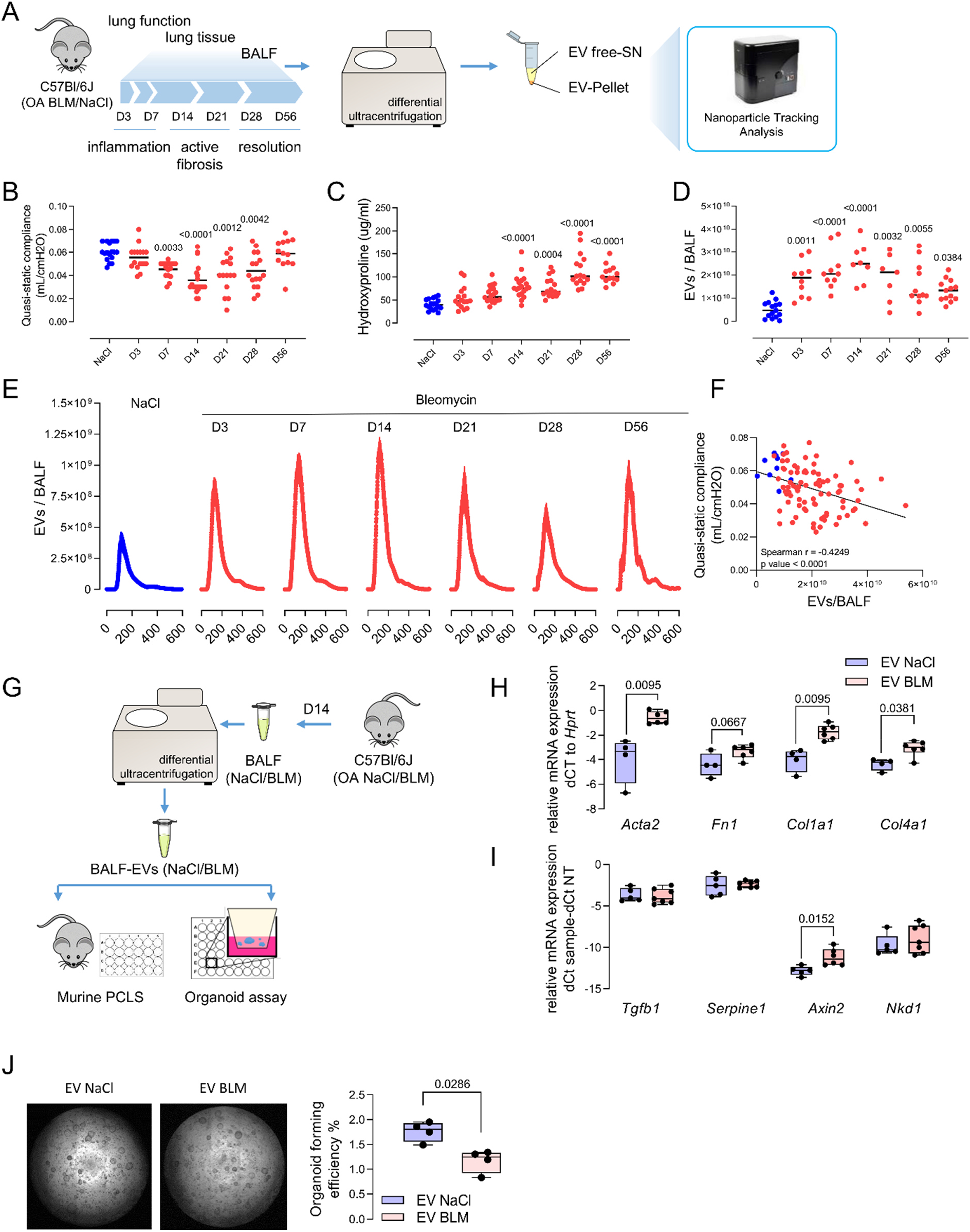
EVs accumulated in lung fibrosis, initiate lung remodeling and impaired alveolar epithelial cell function. **A)** C57Bl/6J mice have been exposed with oro-tracheal bleomycin or NaCl/PBS as control. Lung function was assessed and lung tissue and BALF were collected over the indicated time course. EVs were isolated from BALF and characterized. **B)** Quasi-static compliance and **C)** hydroxyproline level of the corresponding experiment are shown. **D)** BALF-EVs have been numbered by Nanosight and are expressed as the number of particles per BALF. **E)** EV quantification according to particle diameters (expressed in nm) in each time point after bleomycin exposure. **F)** Correlation between EV number and quasi-static compliance is depicted. B-F) Each point corresponds to a mouse (n=5-8 for NaCl groups, n=13-20 for BLM groups). **G)** BALF-EVs have been isolated from mice with pulmonary fibrosis (14 days post-bleomycin) or control mice and used for functional assays. **H, I)** PCLS from normal C57Bl/6J have been cultured with the above-mentioned BALF-EVs. After seven days, the expression of fibrosis-related genes has been assessed by qPCR. Data are representative of PCLS from individual mice exposed to control- (n=4 PCLS) or fibrotic-EVs (n=6 PCLS). Gene expression was normalized to *Hprt* expression. **J)** Murine EpCAM positive cells and CCL fibroblasts in matrigel have been exposed to BALF-EVs for 14 days. Representative images (left panel) and quantification (right panel, n=4 control EVs, n=4 fibrotic EVs) of the organoid formation efficiency. Statistical analysis by Kruskal-Wallis followed by Dunn’s multiple comparisons tests (B-D), Spearman correlation test (F) or non-parametric Mann-Whitney (H-J). p values are indicated for each comparison.

We first aimed to establish the functional relevance of EVs for fibrogenesis in the lung. To this end, EVs were isolated from mice exposed to bleomycin for 14 days (called fibrotic EVs from hereon) or saline control (called control EVs from hereon). The functional properties of these vesicles were tested assessing profibrotic markers in multicellular precision-cut lung slices (PCLS) and alveolar epithelial type (AT) 2 stem cell based organoid assays (figure 1G). PCLS generated from naïve C57Bl/6 mice were treated with fibrotic or control EVs, respectively, and fibrotic markers were assessed by qPCR.

Notably, fibrotic but not control EVs induced the expression of ECM and fibrosis markers such as α-smooth muscle actin or α-SMA (*Acta2*, p value 0.0095), fibronectin (*Fn1*, p value 0.0667), type 1 collagen (*Col1a1*, p value 0.0095) and type 4 collagen (*Col4a1*, p value 0.0381) in healthy lung tissue *ex vivo* (figure 1H). Moreover, WNT signaling is known to be aberrantly active in pulmonary fibrosis (Konigshoff et al., 2008) and we found that the WNT target gene *Axin2* is significantly upregulated by fibrotic EVs compared with control EVs (figure 1I). Impaired lung epithelial cell function is another hallmark of pulmonary fibrosis (Konigshoff et al., 2009). The progenitor cell function of AT2 stem cells has been shown to be reduced in pulmonary fibrosis (Xie et al., 2021). In line with this, we found significantly less organoids when AT2 cells were exposed to fibrotic EVs compared with control vesicles (figure 1J). These findings strongly support the notion that fibrotic EVs carry cargo that is sufficient to drive fibrosis development.

### Fibrotic EVs have a specific proteomic profile

To identify the cargo responsible for the pro-fibrotic activity of EVs, we comprehensively characterized their proteomic profile. We produced BALF-EVs and corresponding EV-free BALF fractions from fibrotic and healthy mouse lungs, respectively (figure 2A). After quality control (figure S1A), the samples were subjected to unbiased label-free shot-gun proteomics. This approach identified 1634 proteins overall, including 774 proteins specifically enriched in the EV samples (94 only in BLM-EVs, 389 only in PBS-EVs) and 218 specific to the EV-free samples (figure S1B). Both, principal component analysis and the heatmap generated after unsupervised hierarchical clustering of the samples based on Pearson correlation revealed significant differences in the proteome profiles amongst the samples (figure 2B, 2C). Importantly, we observed enrichment for proteins classically identified in EVs (as described in vesiclepedia.com) in the vesicular fractions compared to EV-free fractions with proteins like CD9, CD81, Flotillin-1 or TSG101, validating our EV isolation protocol (figure S1C). Moreover, fibrotic EVs showed an enrichment in classic fibrosis markers compared to control EVs (such as Tenascin C, MMP19, and collagens) (figure S1D). Using unsupervised clustering, we identified 7 specific clusters demonstrating distinct protein profiles between control and fibrosis, and remarkably, also between EVs and EV-free samples (figure 2C). These clusters were enriched for specific biological processes based on GO terms (figure 2D). Cluster D illustrated the differences between EVs and EV-free fractions and was composed mainly of proteins important for EV generation and secretion, such as Rab proteins, caveolin and flotillins. More importantly, cluster B contained 107 EV proteins which were specifically enriched in fibrotic EVs compared to normal EVs and were not changed in the corresponding EV-free supernatant (EV-free SN) fraction (figure 2C). This cluster included several proteins involved in fibrosis, cell adhesion or extracellular matrix organization (figure 2D). We confirmed the specific enrichment in fibrotic EVs of four proteins identified by our unbiased proteomic approach (figure 2E, 2F). Among them, Nidogen-1, AGER and EGFR were previously linked to the mechanisms of fibrogenesis (Englert et al., 2008; Tzouvelekis et al., 2013) and some of which have been found to be secreted by cells via EVs (Mao et al., 2020; Patterson et al., 2018; Zhang et al., 2017). By further characterizing cluster B proteins using a GO-based annotation, we found that fibrotic EV cargo was linked to GO-terms such as biological regulation, response to stimulus, developmental process or cell proliferation (figure S1E). We also observed significant enrichment for fibrosis-relevant GO-terms such as extracellular matrix organization (FDR 3.13E^-7^), cell-substrate adhesion (FDR 1.96E^-9^) and wound healing (FDR 2.28E^-6^) (figure S1F). Analysis of the reactome of these fibrotic EV proteins highlighted pathways such as laminin interactions, MET signaling, cell motility and degradation of the extracellular matrix (figure S1G). Altogether, these bioinformatic analyses suggest that fibrotic EVs are enriched in distinct proteins that are relevant to fibrosis.

**Figure 2.**
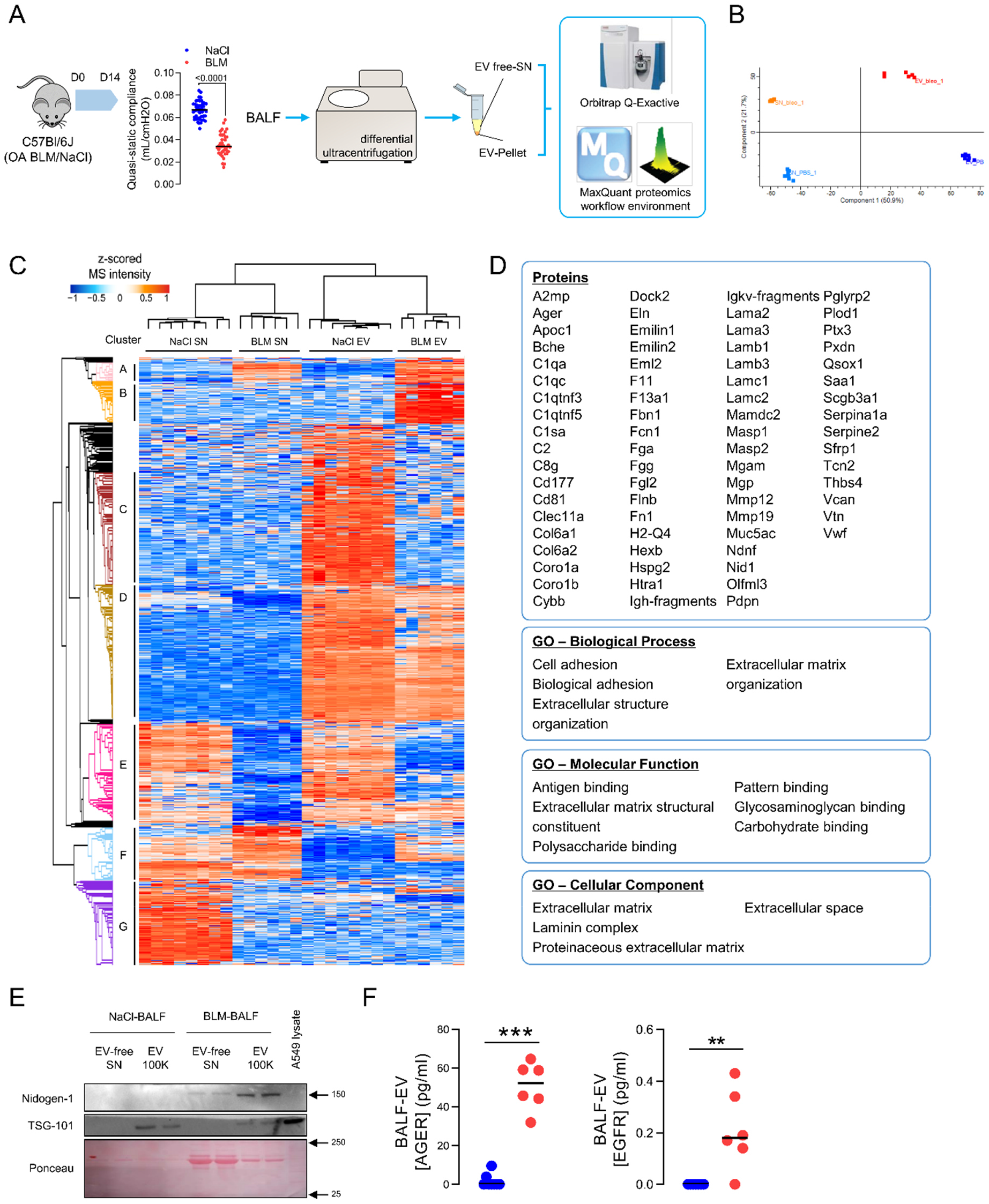
Label-free proteomics identified several proteins specific to fibrotic EVs. **A)** C57Bl/6J mice have been exposed with oro-tracheal bleomycin or NaCl/PBS as control. 14 days post injection, lung function was assessed and lung tissue and BALF were collected. BALF were utilized for EV isolation and vesicular (EV-100K) as well as non-vesicular counterparts (EV-free SN) were subjected to label-free proteomic (n=8 for control, n=6 for bleomycin). **B)** PCA representation of the different samples. **C)** Heatmap of the identified proteins. Color coding corresponds to z-scored MS intensity values after imputation. Based on the unsupervised clustering, proteins were grouped into seven clusters called A to G. **D)** GO analysis of the cluster B. **E)** Levels of Nidogen-1 (western blot) in EVs from bleomycin or control mice are shown. TSG-101 used as protein enriched in EVs and protein content for each sample is shown with ponceau. A549 lysate serves as positive control. Equal (10μg) protein content has been used for the western blot. **F)** AGER and EGFR levels assessed by ELISA on normal (blue) and fibrotic (red) EVs. Data are presented as analyte concentration (pg/ml) normalized to 2×10^8^ vesicles. Statistical analysis by non-parametric Mann-Whitney with **p<0.01, ***p<0.001. SN, EV-free fraction; BLM, Bleo, bleomycin.

### Fibroblasts are a major source of EVs in lung fibrosis

IPF results from an impaired cellular crosstalk involving aberrantly activated epithelial, mesenchymal and immune cells, respectively. We thus aimed to shed light on the key cellular origin(s) of the EVs by annotating the proteins from our proteomic dataset with an already published scRNAseq dataset of the bleomycin model (Strunz et al., 2020). We focused on cluster B and the 107 proteins significantly enriched in fibrotic EVs compared with control vesicles. From these proteins, 47 proteins were linked to a specific cellular annotation, including alveolar epithelial cell, macrophage, and fibroblast subpopulations. Interestingly, we found that the highest number of proteins were linked to fibroblasts (figure 3A). To corroborate this further, we used a single cell RNA sequencing dataset of experimental lung fibrosis (Strunz et al., 2020) to investigate which cell type expressed the highest level of the proteins found in fibrotic EVs. We scored each cell type according to the expression of protein signatures identified in control EVs, fibrotic EVs or in both EV groups together (figure S4). Consequently, our analysis revealed that mesenchymal cells are the main source of a 107 proteins-containing signature specifically enriched in fibrotic EVs (figure 3B). Of note, similar differences were not observed for the expression of control EVs or EV-specific proteins, for which the overall expression remained stable across the cell clusters (figure S4C-D). Moreover, within these stromal cell types, fibroblasts emerged as the major expressers of the fibrotic EV protein signature (figure 3C). We then focused on the fibroblast cell population and ranked the 9 fibroblast-specific proteins of interest according to their expression levels. This analysis revealed that from the originally identified fibrotic EV protein signature, *Serpine2*, *Sfrp1* and *Eln* were the three highest-ranked genes/proteins expressed in fibroblasts (figure 3D).

**Figure 3.**
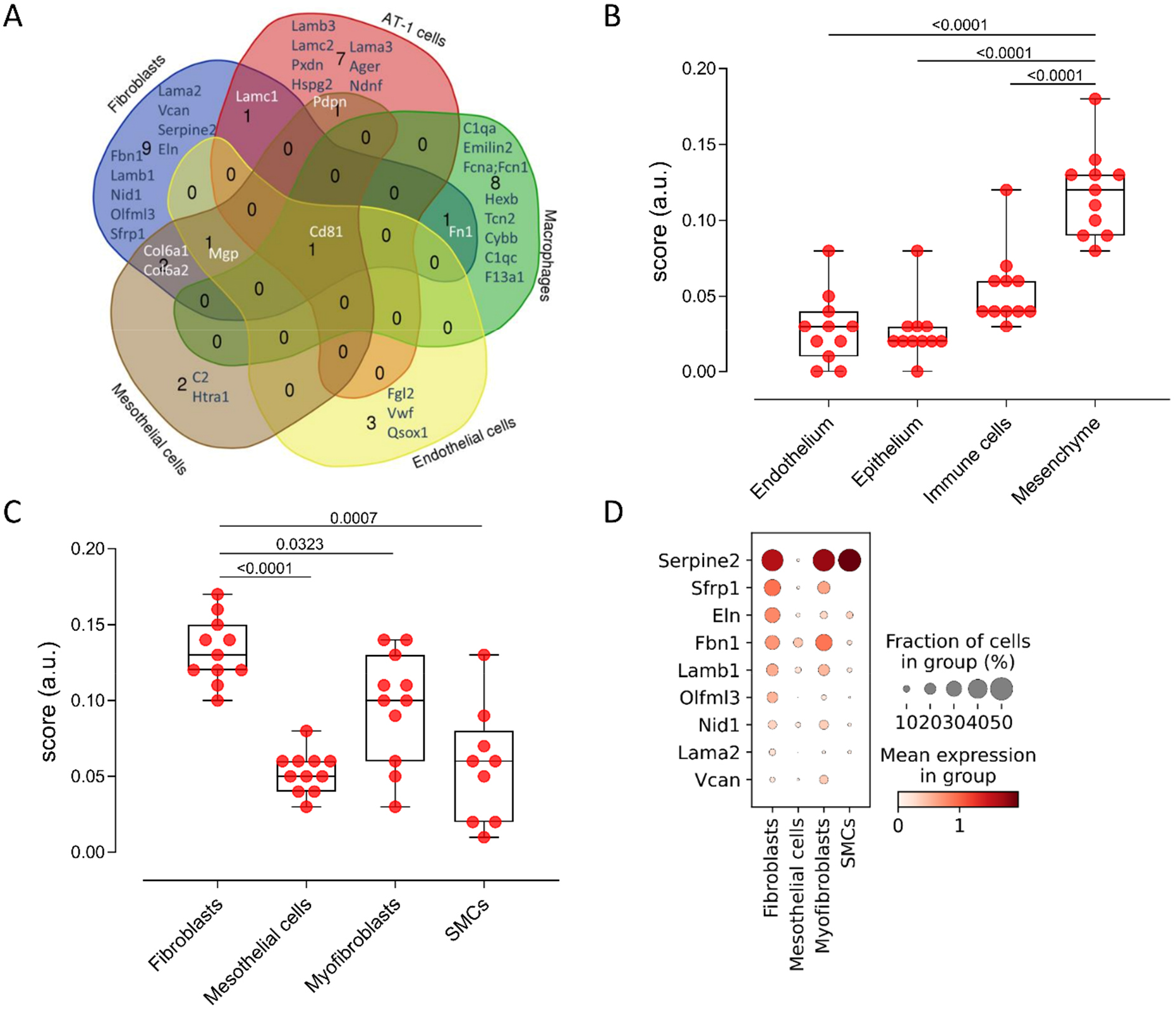
Fibroblasts are a major source of EVs during fibrosis. **A)** Venn diagram depicting the cellular origin of the bleomycin BALF-EV proteins. **B-C)** Scoring of a scRNAseq database (GSE141259) for the mean expression of the 107 proteins identified in the bleomycin BALF-EVs in the main cellular compartments of the lung (B) as well as in mesenchymal populations (C). **D)** Top proteins expressed in fibroblasts, among the proteins identified in bleomycin-BALF-EV. Statistical analysis by non-parametric Mann-Whitney. p values indicated.

### SFRP1 is overexpressed and secreted by fibroblasts during pulmonary fibrosis

From these three top hits, we focused our analysis on SFRP1, a known regulator of WNT signaling (Burgy and Konigshoff, 2018), which is altered in fibrosis (De Langhe et al., 2014; Matsuyama et al., 2014). We observed by immunofluorescence in lung sections that SFRP1 expression is restricted to IPF tissue, particularly to the areas of dense fibrosis characterized by α-SMA staining (figure 4A).

**Figure 4.**
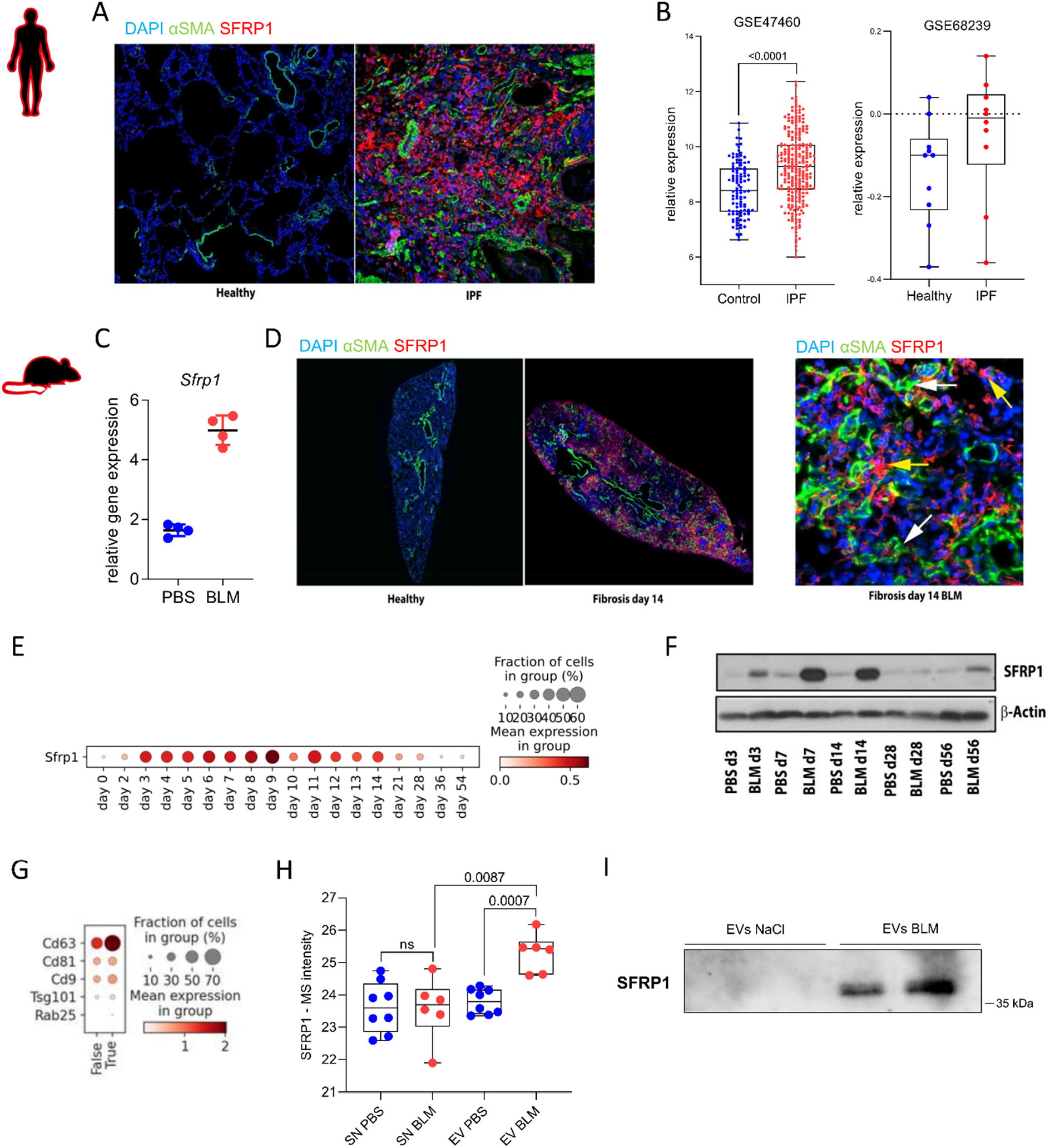
Fibroblast-derived SFRP1 is increased and enriched in EVs during pulmonary fibrosis. **A)** Immunofluorescence staining for aSMA (green) and SFRP1 (red) in FFPE lung parenchyma sections from patients with IPF or healthy donors. DAPI stains nuclei (blue). Representative images are shown. **B)** Gene expression of *SFRP1* in lung tissue from patients with IPF and healthy donors. Data extracted from GSE47460 and GSE68239. **C)** Gene expression of *Sfrp1* in mice with bleomycin-induced lung fibrosis or control (PBS) mice. Data extracted from Schiller *et al*., 2015. **D)** Immunofluorescence staining for aSMA (green) and SFRP1 (red) in FFPE lung sections from mice challenged with bleomycin (day 14) or PBS as control (left panel). Representative observation of a fibrotic area of a mouse lung 14 days post-bleomycin. Yellow arrows highlight SFRP1-expressing cells while white arrows indicate cells positive for αSMA. DAPI stains nuclei (blue). **E-F)** Expression of SFRP1 in lung tissue from mice exposed to bleomycin at different time points at the transcriptomic (E, data from GSE141259) or proteomic level (F). **G)** Analysis of a scRNAseq database (GSE141259) for the expression of EV machinery in cells expressing (true) or not (false) *Sfrp1*. **H)** SFRP1 relative MS intensity in the different fractions analyzed by proteomic as described in figure 1. **I)** Western blot showing SFRP1 accumulation on independent normal and fibrotic EVs (n=2/group). Statistical analysis by parametric unpaired t test (B). p value is indicated.

Using published microarray datasets, we confirmed that *SFRP1* is overexpressed in the lung of patients with IPF compared with controls (figure 4B). In the mouse bleomycin model, *Sfrp1* was overexpressed at day 14 after challenge (figure 4C, 4D). In the lung of mice challenged with bleomycin, we noticed that SFRP1+ cells are localized in areas of active fibrosis containing α-SMA positive cells (figure 4D). SFRP1 expression increased starting 3 days post-bleomycin until day 14 compared with correspondent saline controls, before decreasing and returning back to baseline at day 28 (figure 4E, 4F). Transcript analysis revealed that among the five members of the SFRP1 family, *SFRP1* is the highest expressed form in murine or human normal lung fibroblasts (figure S5). In addition, *Sfrp1* expressing fibroblasts have enhanced expression of the machinery of EV/exosome biogenesis (figure 4G). In line with our proteomic approach, we confirmed increased SFRP1 in EVs isolated from mice with pulmonary fibrosis (figure 4H, 4I). *In vitro*, we found SFRP1 proteins in EVs secreted from fibroblasts, and not in the EV-free fractions of the cell supernatant (figure S6).

### SFRP1 inhibition hampers the pro-fibrotic activities of EVs

We next aimed to determine whether SFRP1 can be a target that attenuates the action of fibrotic EVs. As shown earlier, SFRP1 is a protein produced by fibroblasts during pulmonary fibrosis. In this set of experiments, we used primary lung fibroblasts isolated from *Sfrp1* deficient (*Sfrp1^-/-^*) mice or control (*Sfrp1^+/+^*) mice (figure 5A). EVs were concentrated from the conditioned media of *Sfrp1^-/-^* and *Sfrp1^+/+^* fibroblasts and injected intra-tracheally into mice previously challenged with bleomycin, based on a previously published protocol (Parimon et al., 2019). EVs were administered repeatedly starting day 8 post-bleomycin as illustrated (figure 5A). We observed that EVs from WT fibroblasts significantly aggravate bleomycin-induced fibrosis while the mice receiving intra-tracheal EVs from Sfrp1*^-/-^* fibroblasts exhibited significantly less fibrosis after 21 days (figure 5B). Importantly, mice exposed to EVs from Sfrp1*^-/-^* fibroblasts had significantly less collagen accumulation within the lung compared with mice receiving EVs from WT fibroblasts (figure 5C).

**Figure 5.**
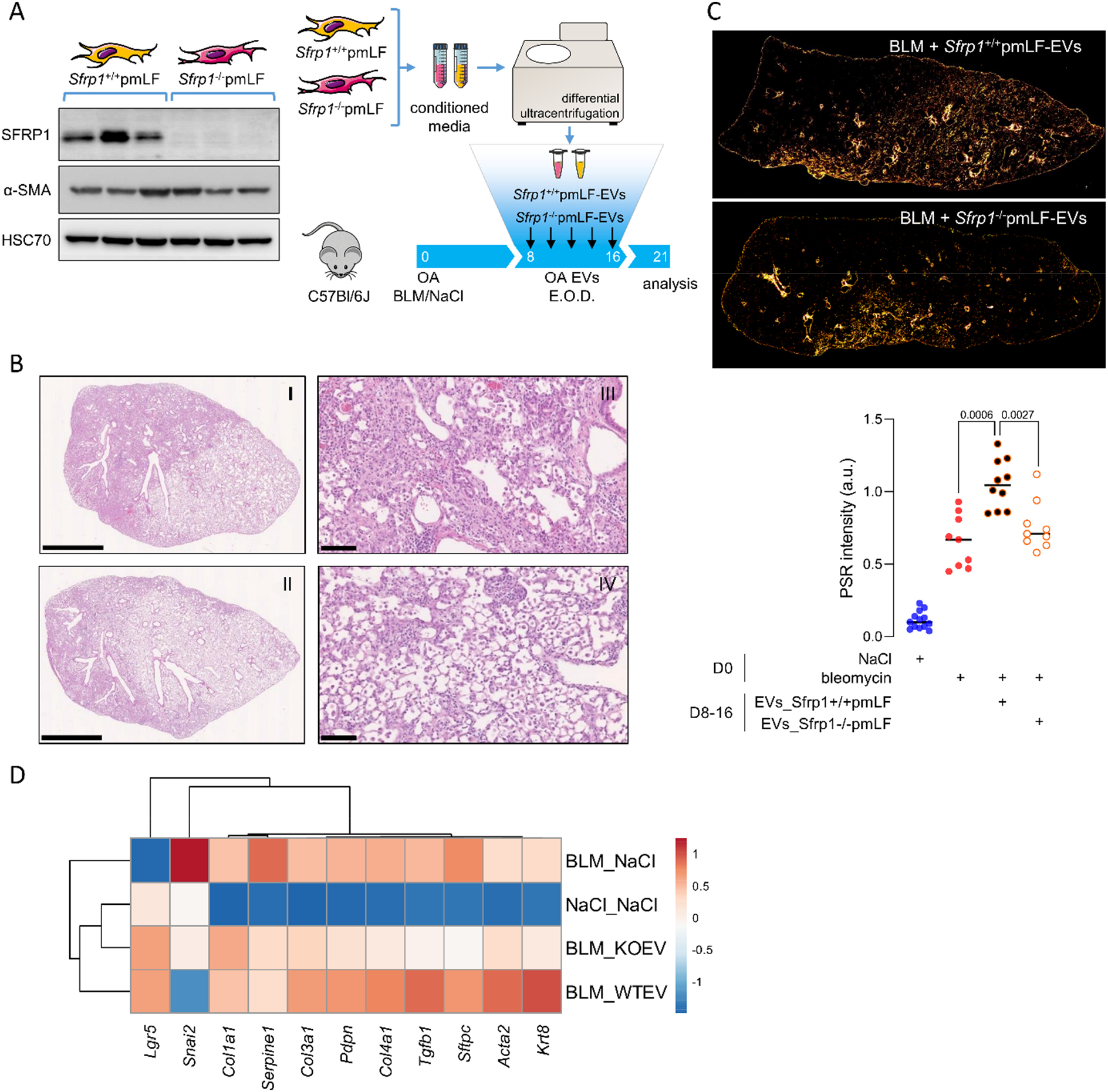
*Sfrp1* deficiency in fibroblast-derived EVs attenuates lung fibrosis in vivo. **A)** General outline of the experiment. EVs have been isolated from the conditioned media of primary mouse lung fibroblasts coming from mice with the genetic deletion of Sfrp1 (Sfrp1-/-) or wild-type control (Sfrp1+/+). SFRP1 expression was confirmed by western blot (left panel). Isolated vesicles were injected intra-tracheally in mice previously challenged with bleomycin or control NaCl. **B)** Representative histology of the lung of the above described mice at D21 post bleomycin exposure. Panels I and III are representative of the “BLM+WT_EVs” and II and IV representatives of the “BLM+Sfrp1-/-_EVs”. Scale bars indicate 2.5 μm (I, II) or 100 μm (III, IV). **C)** Collagen quantification on FFPE lung sections stained with picro-sirius red and visualized under polarized light. Representative observation (n=10 for BLM+WT_EVs and n= BLM+Sfrp1-/-_EVs, upper panel) and quantification (lower panel) are shown. **D)** RNA-seq was performed on whole lung tissue of the above described mice at D21. Relative expression (z-score) of fibrosis-related genes shown in the heatmap. Statistical analysis by non-parametric Mann-Whitney. Each point represents one mouse. P values are indicated for each comparison.

We performed a transcriptomic analysis by subjecting the lung tissue of these mice to RNA-sequencing. Here, we found that in mice challenged with both bleomycin and *Sfrp1^-/-^* EVs (BLM_KOEV) the expression of collagens (*Cola1a, Col3a1, Col4a1*) along with prominent (myo)fibroblast activation markers *Acta2* and *Tgfb1* were reduced compared to *Sfrp1^+/+^* EVs (BLM_WTEV) (figure 5D). These findings were in accordance with the histological analysis demonstrating an amelioration of lung-fibrosis with *Sfrp1^-/-^* EVs exposure. Moreover, we further discovered that the expression of keratin 8 (*Krt8*), a recently described marker of aberrantly activated AT2 cells, was increased in mice exposed to vesicles from WT fibroblasts compared to *Sfrp1^-/-^* fibroblasts EVs.

## Discussion

In the past years, increased evidence has emerged showing impaired cellular crosstalk during fibrosis (Distler et al., 2019; Kurche et al., 2022). Consequently, the study of EVs including exosomes has gained significant attention in the field (Brigstock, 2021). These secreted vesicles are effective mediators of cell-to-cell communication and participate in physiological and also pathological processes. Due to their accumulation in body fluids and their ability to participate in disease mechanisms, EVs are regarded as interesting targets to develop both novel therapies and also diagnosis/prognostic tools (Soekmadji et al., 2020). We and others demonstrated the crucial role of EVs during pulmonary fibrosis (Martin-Medina et al., 2018; Njock et al., 2019; Parimon et al., 2019; Santos-Alvarez et al., 2022). These vesicles accumulate within the lungs of patients with IPF and can be collected in several body fluids such as BALF, sputum or urine (Elliot et al., 2022; Martin-Medina et al., 2018; Njock et al., 2019). During fibrosis, EVs exacerbate major mechanisms of disease such as proliferation and activation of fibroblasts (Elliot et al., 2022; Martin-Medina et al., 2018) or AT2 cells (Parimon et al., 2019). Besides, EVs may also have anti-fibrotic properties via the inhibition of TGF-β signaling (Guiot et al., 2020). These functions rely on the transfer of specific cargoes from one cell to another (van Niel et al., 2018). Considerable efforts have been undertaken to investigate EV’s cargo and their corresponding biological function. In lung fibrosis several unbiased approaches identified specific cargo signatures of fibrotic vesicles mainly for microRNAs (Njock et al., 2019; Parimon et al., 2019; Santos-Alvarez et al., 2022). In particular, increasing levels of specific miRNAs transported by extracellular vesicles are linked to the activation of pathological signaling pathways and correlate with declined lung function in patients (Njock et al., 2019; Parimon et al., 2019). However, our knowledge about EVs’ protein cargo is still limited (Burgy et al., 2020). In addition, the heterogeneity of these vesicles adds to the difficulty of achieving a deeper characterization of fibrotic EVs. Identifying the cells secreting vesicles during fibrosis is also a key prerequisite to mechanistically understand the biological function of fibrotic EVs. Here, we uncover a key mediator of pro-fibrotic EVs. We demonstrate that 1) secreted EVs exhibit a distinct proteome profile in pulmonary fibrosis, 2) fibrotic EVs alone trigger ECM accumulation and reduce AT2 stem cell function, and 3) the presence of SFRP1 in fibrotic EVs is required, at least partially, for the pro-fibrotic effect of these vesicles.

In the present study, we aimed to deeply characterize the specific proteome of EVs during fibrosis. For a meaningful analysis of extracellular vesicles and their functional effects, the methodology of vesicle concentration is crucial (Thery et al., 2018). We applied a well-established and widely used protocol based on differential ultracentrifugation (Coughlan et al., 2020; Martin-Medina et al., 2018). As an additional separation step, we implemented a size exclusion chromatography before mass spectrometry characterization of our isolated EVs, as previously described (Kavanagh et al., 2017). This step served to get rid of components smaller than 30 nm such as protein aggregates. Next to state-of-the-art assessment via NTA and electron microscopy (Thery et al., 2018), our successful EV fractionation was further confirmed by our proteomic approach. We observed the enrichment, in the EV samples, of canonical EV proteins (TOP100 listed in Vesiclepedia) such as CD9, CD81, Flotillin-1 etc. This validates our approach which leads to the first description of the proteome of EVs during homeostasis and pulmonary fibrosis. While our data are in line with previously reported EV proteomes from patients with IPF (Shaba et al., 2021), we made a step further and identified the proteins transported by fibrotic EVs specifically (*i.e*. not on control vesicles, and not on EV-free fraction). To this end, we used the EV-free fractions to filter the proteins enriched in EV-free fractions of fibrotic samples over respective controls. The relevance of our approach relies on this double filtering, allowing us to discriminate between proteins secreted during fibrosis, and proteins specifically transported by EVs during the disease. We then confirmed proteins already described as enriched in fibrotic EVs such as fibronectin (Chanda et al., 2019), and identified SFRP1 as a novel protein specific to fibrotic vesicles.

Our findings further corroborate recent reports showing increased EVs in pulmonary fibrosis (Guiot et al., 2020; Kadota et al., 2021; Martin-Medina et al., 2018). However, the cellular source of these secreted vesicles is still largely unknown. Here, we used an unbiased profiling of EVs during fibrosis to shed light on their cellular origin. Indeed, we found distinct protein signatures associated with specific cell types within the lung, indicating that EVs isolated from BALF samples are secreted by immune and structural cells. While we detected proteins expressed by all cell types in non-disease control samples, proteins identified in fibrotic EVs were mainly linked to fibroblasts and alveolar epithelial cells. Our bioinformatic analysis using single cell RNA sequencing datasets identified fibroblasts as the major source of fibrotic EVs. In our seminal work showing increased accumulation of EVs in the lung of patients with IPF, we demonstrated that these vesicles are loaded with WNT5A (Martin-Medina et al., 2018). During IPF, WNT5A is mainly expressed by activated (myo)fibroblasts (Vuga et al., 2009) suggesting that the increase of the EVs seen in IPF patients are due, at least partially, by an increase of vesicle secretion from fibroblasts. Similarly, others have reported that pathological fibroblasts, activated by pro-fibrotic cytokines or made senescent, secrete higher levels of EVs (Chanda et al., 2019; Kang et al., 2020; Lacy et al., 2019). Altogether, this highlights fibroblasts as a major (although not exclusive) source of EVs during pulmonary fibrosis. Of note, epithelial cells have also been described as relevant secretors of fibrotic EVs (Parimon et al., 2019). Beyond EV secretion, the nature of EVs’ recipient cells remains elusive and further research is needed to decipher the cross talk mediated by fibrotic EVs by auto- or paracrine signaling.

A growing body of literature indicates that EVs exhibit pro-fibrotic roles. We previously showed that EVs from fibroblasts activated by TGF-β1 induced fibroblast proliferation in a WNT5A dependent manner (Martin-Medina et al., 2018). Notably, the addition of fibrotic EVs enhanced collagen accumulation within the lung *in vivo* (Parimon et al., 2019). Several other studies report a profibrotic effect of EVs, mediated by protein cargo, during pulmonary fibrosis (Kang et al., 2020; Lacy et al., 2019). Senescent fibroblasts secrete higher numbers of EVs coated with fibronectin on their surface (Chanda et al., 2019) and thus promote the invasive capacity of fibroblasts via the interaction of fibronectin and α5β1 integrins. In line, lung fibroblasts isolated from patients with IPF secrete EVs which in turn induce senescence in lung epithelial cells *in vitro* (Kadota et al., 2020). This effect seems to be mediated by the transfer of microRNAs leading to mitochondria destabilization within the recipient cells. Altogether, the data presented in our study are in agreement with current literature indicating that EVs aggravate fibrosis. Here, we confirmed that fibrotic EVs are functional and induced the expression of fibrosis-related genes in normal lung tissue *in vitro*. We found this effect to be mediated by perturbation of the developmental WNT signaling pathway linked to cellular senescence and fibroblast differentiation according to previous reports (Lehmann et al., 2020). Indeed, we have previously shown that the chronic activation of WNT/β-catenin signaling results in a robust induction of senescence in AT2 cells, which thus exhibit decreased progenitor cell potential (Lehmann et al., 2020). This supports our observations that fibrotic EVs promote fibrotic-like changes, most likely by impairing the regenerative properties of AT2 cells.

We found fibroblast-derived SFRP1 to be a central protein cargo of EV mediated pulmonary fibrosis. The family of soluble FRP (SFRP) is constituted of five members which can bind WNT ligands, and thus hamper their activities. Our screening identified SFRP1 as a protein specifically enriched in fibrotic EVs, suggesting that their pro-fibrotic properties are mediated by this soluble receptor. Indeed, SFRP1 is secreted by senescent cells as part of the senescence-associated secretory phenotype and can induce paracrine senescence, a major hallmark of pulmonary fibrosis (Elzi et al., 2012). *Sfrp1* deficient mice exhibit enhanced kidney fibrosis after unilateral ureteral obstruction compared with wildtype mice (Matsuyama et al., 2014). In mice challenged with intratracheal bleomycin, the whole-body deletion of *Sfrp1* does not alter fibrosis development (De Langhe et al., 2014). However, we found here that EVs secreted from fibroblasts (carrying SFRP1) exacerbated fibrosis and this effect was abolished with vesicles from *Sfrp1^-/-^* fibroblasts. These conflicting results might be explained in several ways: firstly, SFRPs constitute a family of proteins which have redundant roles and the study on kidney fibrosis suggests that SFRP1 might have a different organ-specific role. Secondly, the *Sfrp1* deficient mice used were whole body knock-out animals. Whether non-vesicular SFRP1 may exhibit specific functions relevant to wound healing and tissue fibrosis is an open question which remains to be answered. Although additional research is needed to fully understand the biology of SFRP1 during fibrosis, our findings suggest the relevance of targeting the EV-linked form of SFRP1 to counteract fibrosis development.

In parallel, some EV subpopulations are likely to have pro-repair functions (Kadota et al., 2021). EVs concentrated from the culture media of bronchial epithelial cells impede both β-catenin dependant and independant WNT signaling pathways, resulting in lowered myofibroblast activation and decreased cellular senescence in epithelial cells. This underlines that EVs are heterogenous within the lung and while some vesicles exacerbate fibrosis mechanisms, others seem to be secreted as an attempt to drive tissue regeneration. Our recent work on alveolar regeneration identified Krt8+ epithelial cells, which is a transitional state in the physiological AT2 to AT1 differentiation. During fibrosis, these cells aberrantly persist probably due to impaired tissue niche interactions and show pathological features such as senescence (Strunz et al., 2020). In the present study, we found that exposure of SFRP1-loaded EVs increased *Krt8* expression in the lung of mice with pulmonary fibrosis. This effect was abolished when EVs without SFRP1 were administered. This even suggests a therapeutic potential of targeting specific EV subpopulations (*i.e*. SFRP1-EVs in our case) to prevent fibrosis progression. Further research is needed to shed light into vesicle heterogeneity in physiological and fibrotic processes. The identification of markers for specific EV populations will be key to target pathologic vesicles.

EVs are secreted by all cells and are released in almost all body fluids making them highly interesting targets in the field of biomarker discovery (Zhou et al., 2020). A significant challenge in the management of patients with IPF comes from the heterogeneity in disease progression. Novel biomarkers for patient stratification and assessment of therapy efficacy are urgently needed. In this context, EVs have gained attention in the IPF field. Unbiased approaches uncovered key differences in the miRNA cargo of EVs isolated from patients with IPF with distinct profile when comparing lung tissue-, BALF- and sputum-derived vesicles (Kaur et al., 2021; Njock et al., 2019). Of note, some of these miRNAs transported by EVs correlate with altered lung function. Similarly, specific proteins have been identified in EVs from patients with IPF (d’Alessandro et al., 2021; Shaba et al., 2021). The protein cargo of these vesicles is related to fibrosis-relevant processes such as antigen presentation or cytoskeleton remodeling (Shaba et al., 2021). In order to identify potential biomarkers from easily accessible bodily fluids, miRNA and protein have been tracked in vesicles isolated from patients’ blood. Some of the identified cargos (*e.g*. miR-21-5p, tetraspanins etc) correlate with patient survival (d’Alessandro et al., 2021; Makiguchi et al., 2016) even though these promising results need to be confirmed in independent and larger cohorts. Here, we identified SFRP1 as a crucial component of fibrotic EVs. Whether the level of vesicular SFRP1 correlates with fibrosis severity remains an open question which will be addressed in followup studies. In line, EVs were also found in urine of patients with IPF (Elliot et al., 2022). The authors reported a specific miRNA composition but little is known about the proteins transported by these vesicles. It would be of interest to study the presence of SFRP1 in urine EVs to test its potential as a non-invasive biomarker for fibrosis.

In summary, we report here for the first time that EVs secreted by fibrotic cells or tissue are able to drive a pathologic response in healthy tissue. During pulmonary fibrosis, these disease-relevant vesicles are mainly secreted by fibroblasts within the lung and carry specific proteins such as SFRP1. Our study also brings a proof of concept showing the potential of inhibiting an EV-linked mediator to counteract fibrosis exacerbation. Further research is required to test the potential of targeting EV-transported mediators to counteract fibrosis development.

## Acknowledgements

The authors are very grateful to all members of the Königshoff laboratory for stimulating and fruitful discussions. We are thankful to Kristina Hatakka, Anastasia van den Berg, Kathrin Hafner, Clemence Stoeckel-Linossier, Sabrina Loriod, Marisa Neumann and Christine Hollauer for excellent technical assistance. We thank the animal facility and the ImaFlow core facility in Dijon for their invaluable help. We wish to thank all patients and their families who participated in this study. We gratefully acknowledge the provision of human biomaterial and clinical data from the CPC-M bioArchive and its partners at the Asklepios Biobank Gauting, the LMU Hospital and the Ludwig-Maximilians-Universität München.

## Author contributions

O.B., B.B.L., A.S., D.S. and M.Le. designed and performed experiments, analyzed data and prepared figures; C.H.M. performed mass spectrometry experiments and analysis as well as integrative analysis of proteome data with scRNAseq data and prepared figures; O.B., M.Le., G.B. and M.K. designed experiments and oversaw all data analysis; H.B.S. supervised proteomic data analysis and experimental design; C.G. contributed to nanoparticles tracking experiments and analysis; T.P. and P.C. provided technical assistance and important intellectual content; M.Li. and A. H. collected and provided human tissue samples; M.M, A.Ö.Y. and O.E. analyzed and interpreted results and brought important intellectual content; O.B., A.Ö.Y., H.B.S., G.B. and M.K. provided resources and funding. O.B., M.Le., G.B. and M.K. drafted the manuscript. All authors have critically revised the manuscript. All authors have read, reviewed and approved the final manuscript as submitted to take public responsibility for it.

## Supplementary material

**Figure S1.**
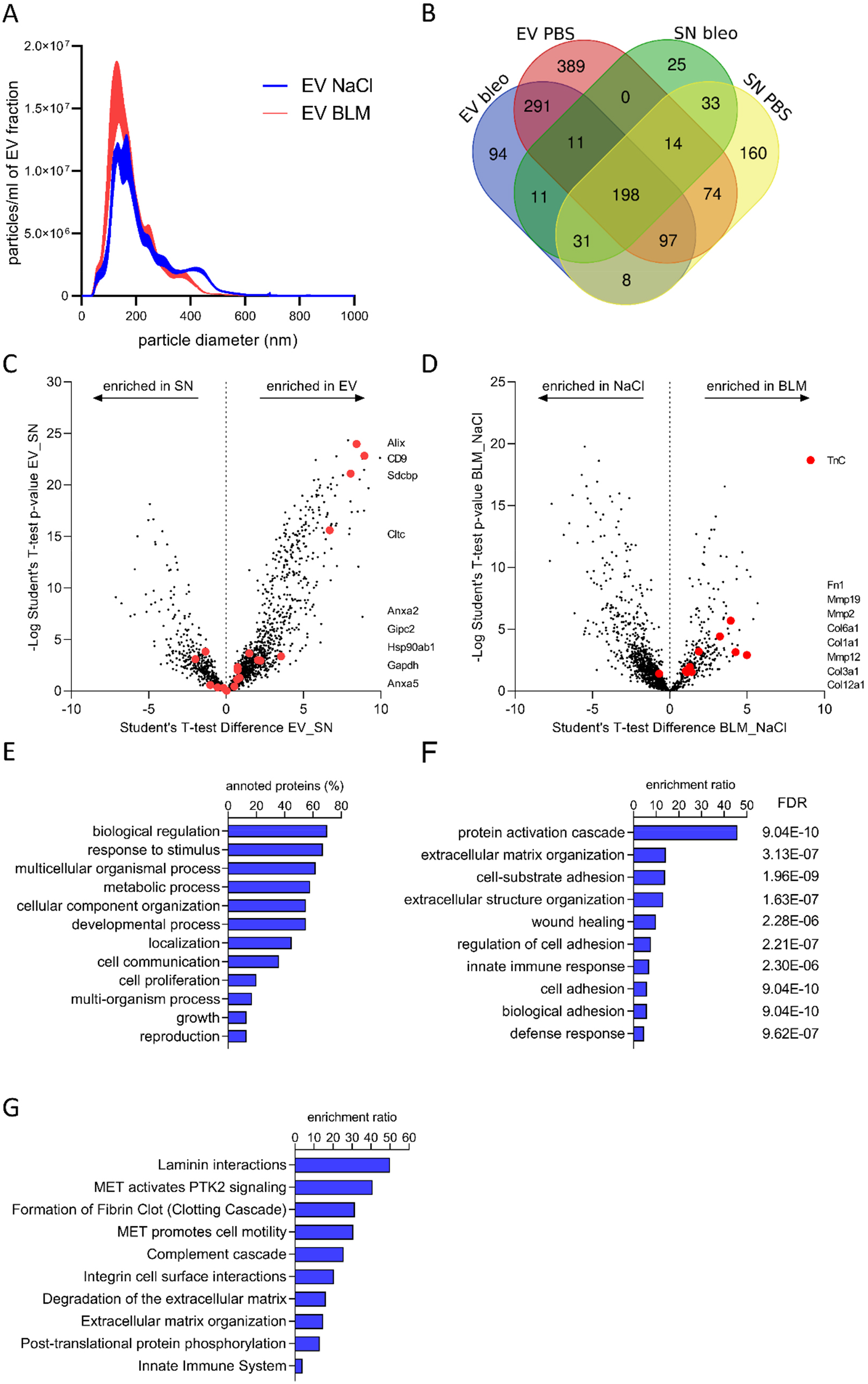
Specific set of proteins are enriched in fibrotic EVs. **A)** Nanoparticle tracking analysis of the EV samples used as input for the proteomic. **B)** Venn diagram showing the number of proteins identified in each fraction or overlapping over several fractions. **C, D)** Volcano plot depicting the proteins differentially regulated in NaCl vs BLM samples (C) or in EV vs EV-free fractions (D). Proteins classically enriched in EVs (Top20 based on Vesiclepedia), or fibrosis relevant proteins are highlighted in red in C and D, respectively. **E-G)** Gene ontology analysis of the 107 POIs for biological process (E), molecular process (F) or reactome study (G).

**Figure S2.**
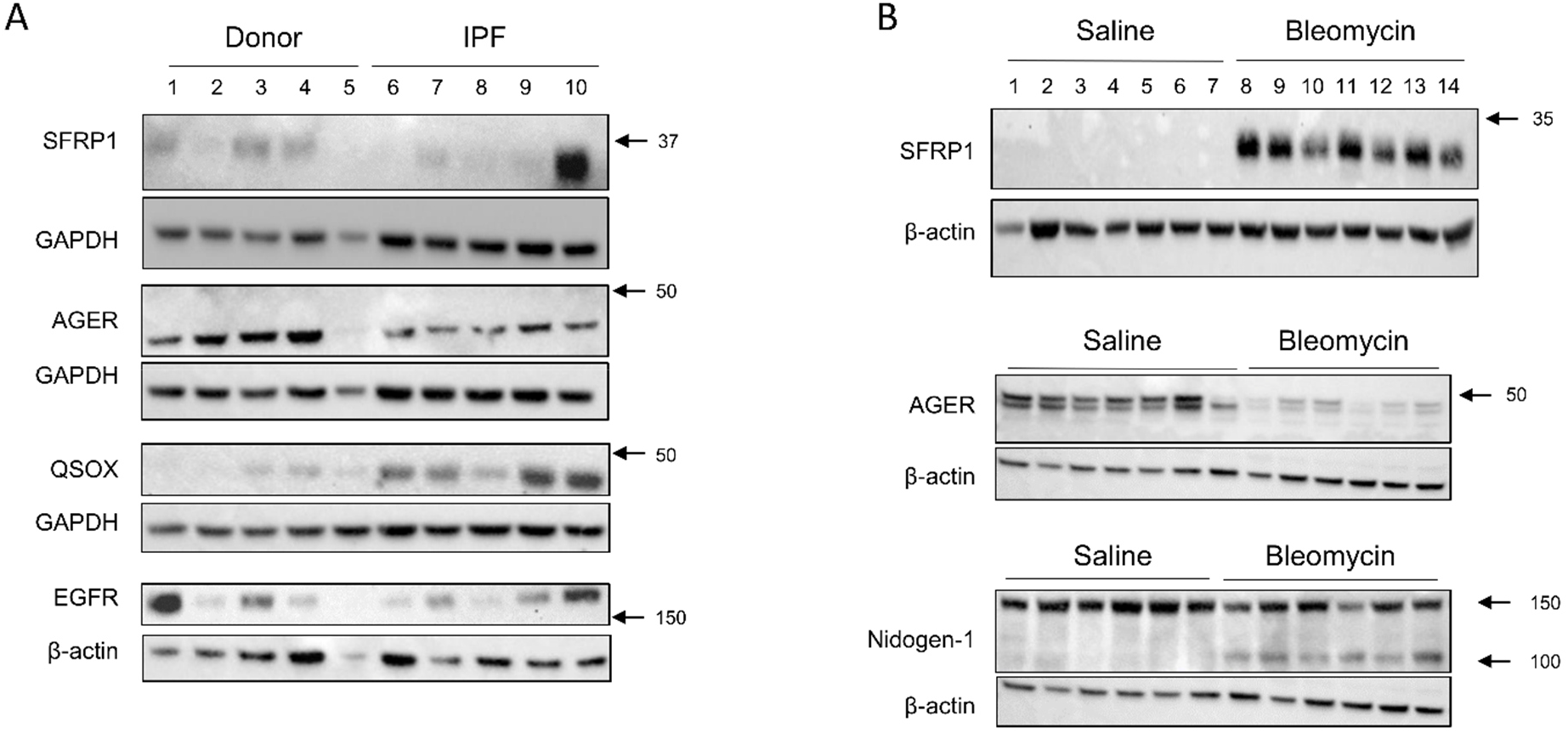
Proteins enriched in fibrotic EVs are dysregulated during pulmonary fibrosis. Expression of the top targets identified by the EV proteomic on tissue from **A)** patients with IPF or healthy donors or **B)** mice with bleomycin-induced pulmonary fibrosis and control mice. GAPDH and β-actin served as loading control.

**Figure S3.**
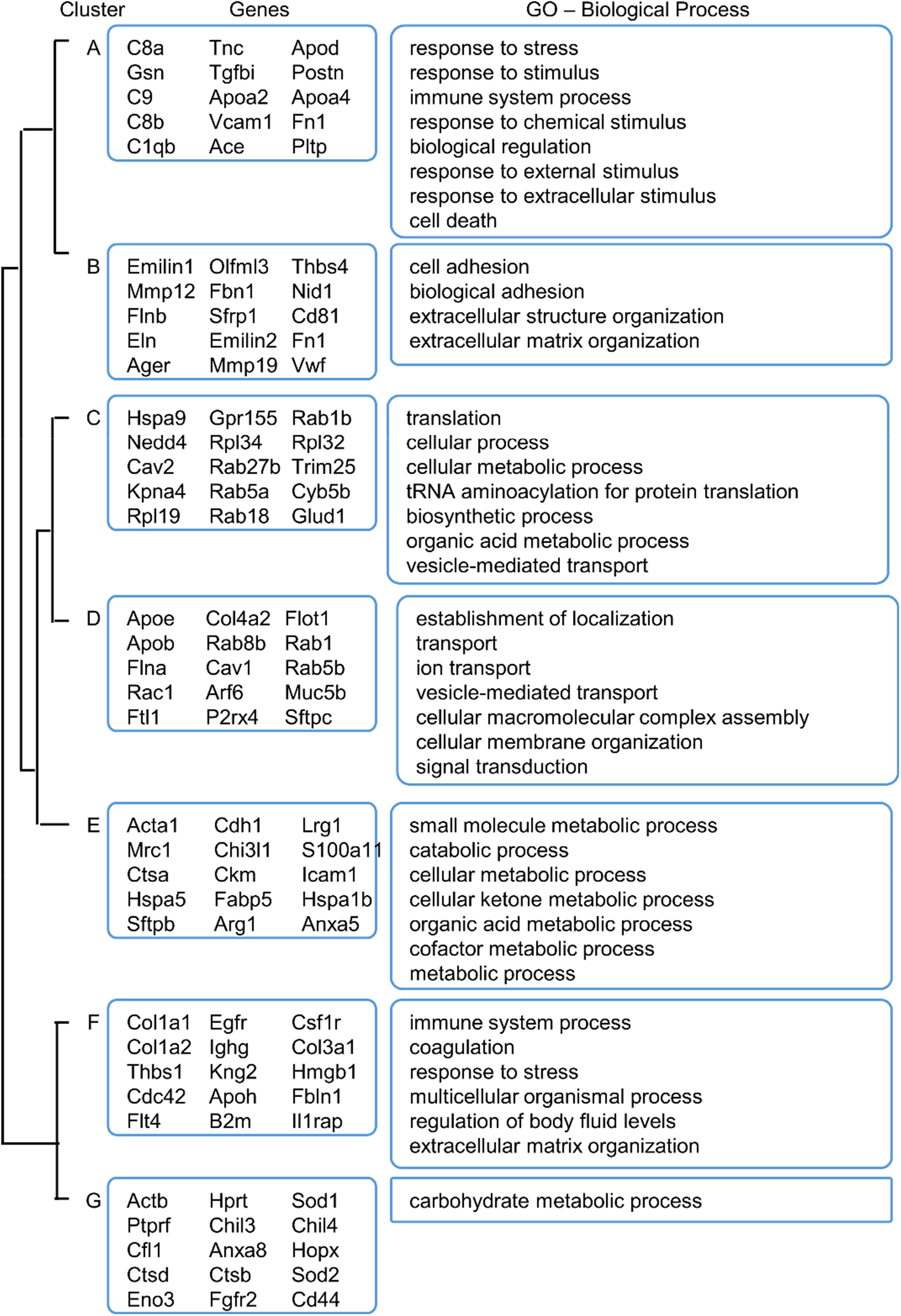
GO annotations of the protein clusters. GO analysis of the clusters identified in figure 2. Top genes are highlighted for each cluster, as well as their GO term for biological process and molecular function.

**Figure S4.**
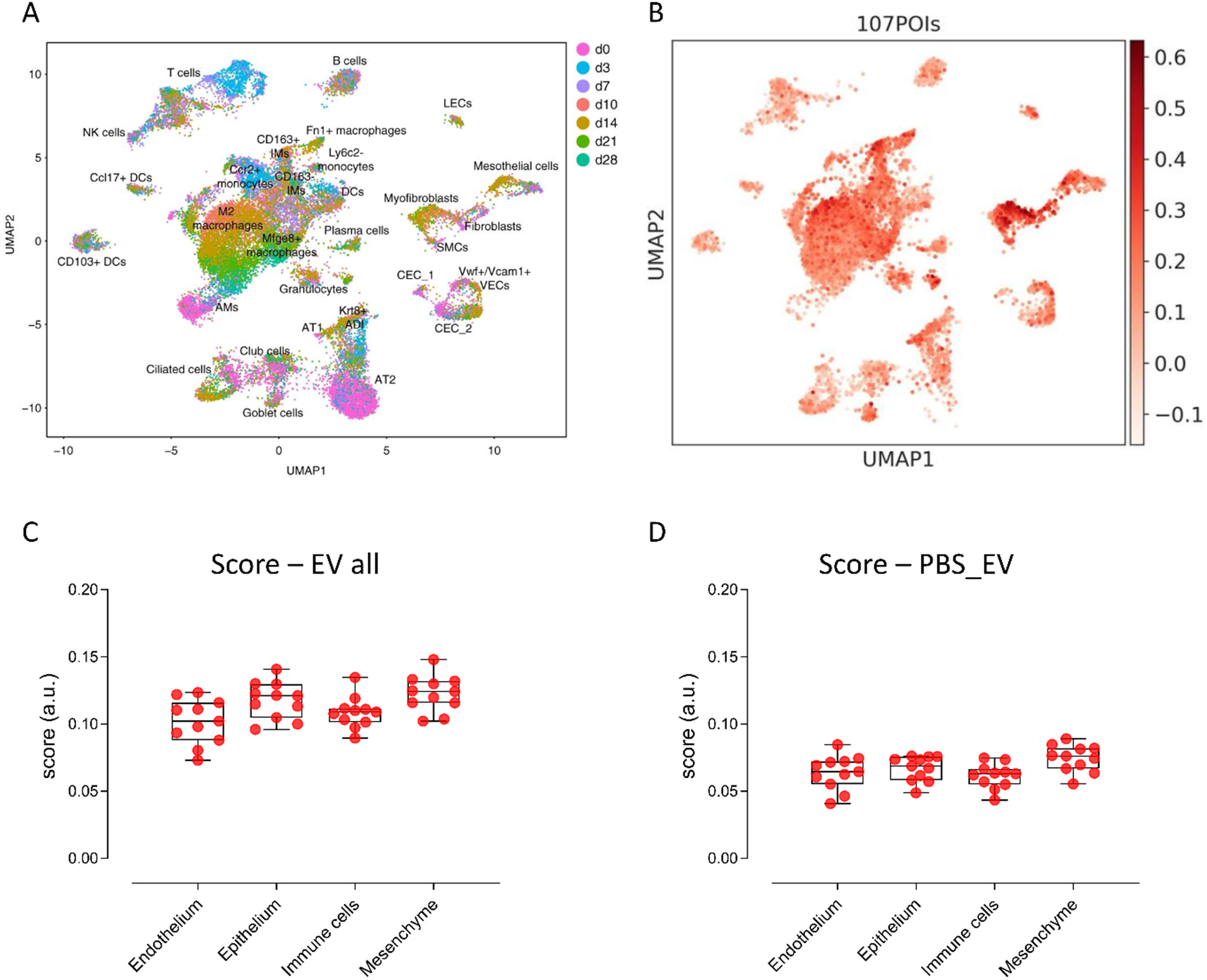
Proteins linked to fibrotic EVs are expressed by mesenchymal cells. **A)** Cell cluster from the scRNAseq dataset analyzed in figure 3 and 4. Data extracted from GSE141259 (Strunz *et al*., 2020). **B)** UMAP representation of the scRNAseq dataset with color coding indicating the scored mean expression of the 107 proteins specific to bleomycin BALF-EVs. **C-D)** Scoring of the scRNAseq dataset for the expression of the proteins identified in the EV samples (C) or specifically in PBS-BALF-EVs (D).

**Figure S5.**
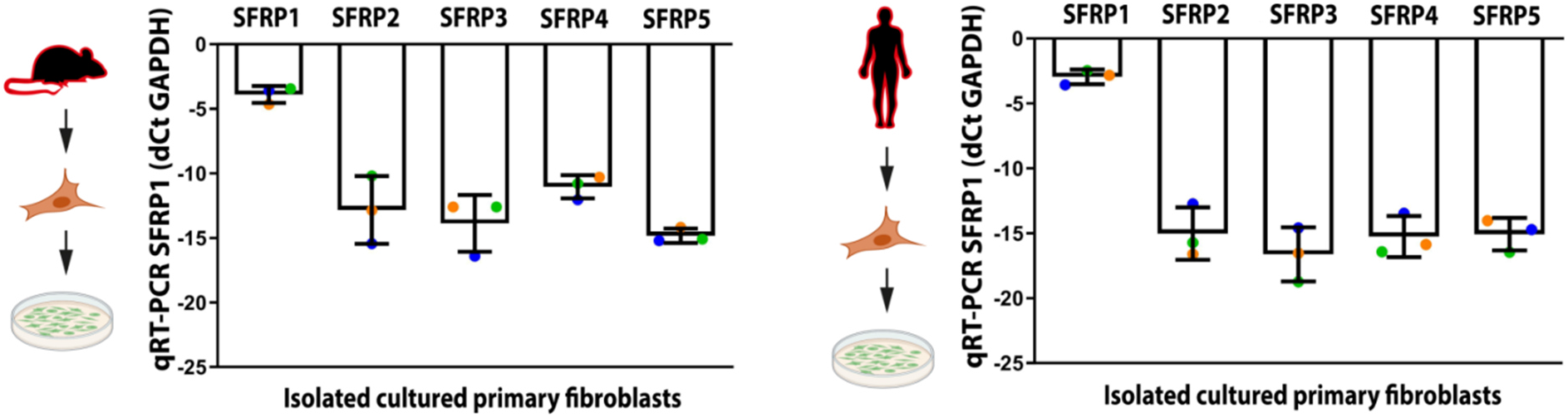
SFRP1 is the SFRP isoform the most expressed in fibroblasts. Primary mouse or human lung fibroblasts have been isolated from healthy animals/donors and cultured *in vitro*. The gene expression of SFRP members was assessed by qPCR. Data representative of three independent experiments.

**Figure S6.**
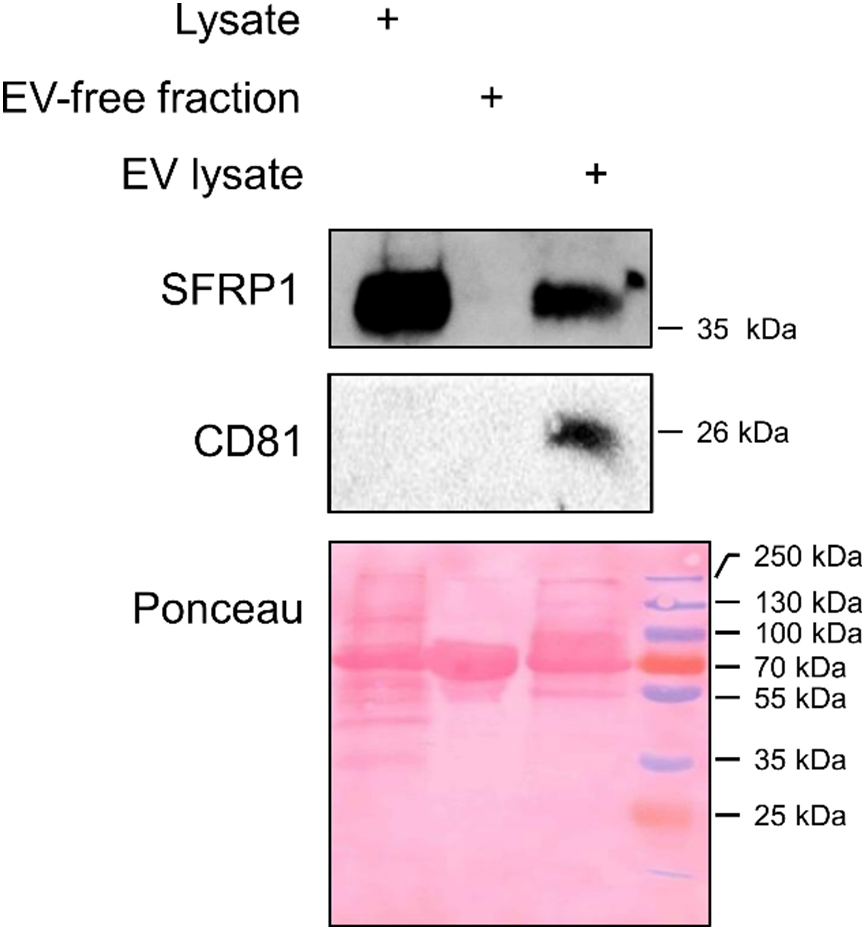
Fibroblasts secrete extracellular vesicles loaded with SFRP1. Primary human lung fibroblasts were cultured and EVs isolated from the cell culture supernatant. SFRP1 expression in the cell lysate, EV-free fraction and EV fractions. CD81 is used as EV-enriched protein and total protein amount is visualized with ponceau.

## References

Bonniaud, P., A. Fabre, N. Frossard, C. Guignabert, M. Inman, W.M. Kuebler, T. Maes, W. Shi, M. Stampfli, S. Uhlig, E. White, M. Witzenrath, P.S. Bellaye, B. Crestani, O. Eickelberg, H. Fehrenbach, A. Guenther, G. Jenkins, G. Joos, A. Magnan, B. Maitre, U.A. Maus, P. Reinhold, J.H.J. Vernooy, L. Richeldi, and M. Kolb. 2018. Optimising experimental research in respiratory diseases: an ERS statement. Eur Respir J 51:

Brigstock, D.R. 2021. Extracellular Vesicles in Organ Fibrosis: Mechanisms, Therapies, and Diagnostics. Cells 10:

Burgy, O., P.S. Bellaye, G. Beltramo, F. Goirand, and P. Bonniaud. 2020. Pathogenesis of fibrosis in interstitial lung disease. Curr Opin Pulm Med 26:429–435.

Burgy, O., and M. Konigshoff. 2018. The WNT signaling pathways in wound healing and fibrosis. Matrix Biol 68-69:67–80.

Chanda, D., E. Otoupalova, K.P. Hough, M.L. Locy, K. Bernard, J.S. Deshane, R.D. Sanderson, J.A. Mobley, and V.J. Thannickal. 2019. Fibronectin on the Surface of Extracellular Vesicles Mediates Fibroblast Invasion. Am J Respir Cell Mol Biol 60:279–288.

Coughlan, C., K.D. Bruce, O. Burgy, T.D. Boyd, C.R. Michel, J.E. Garcia-Perez, V. Adame, P. Anton, B.M. Bettcher, H.J. Chial, M. Konigshoff, E.W.Y. Hsieh, M. Graner, and H. Potter. 2020. Exosome Isolation by Ultracentrifugation and Precipitation and Techniques for Downstream Analyses. Curr Protoc Cell Biol 88:e110.

Cox, J., and M. Mann. 2008. MaxQuant enables high peptide identification rates, individualized p.p.b.-range mass accuracies and proteome-wide protein quantification. Nat Biotechnol 26:1367–1372.

Cox, J., N. Neuhauser, A. Michalski, R.A. Scheltema, J.V. Olsen, and M. Mann. 2011. Andromeda: a peptide search engine integrated into the MaxQuant environment. J Proteome Res 10:1794–1805.

d’Alessandro, M., P. Soccio, L. Bergantini, P. Cameli, G. Scioscia, M.P. Foschino Barbaro, D. Lacedonia, and E. Bargagli. 2021. Extracellular Vesicle Surface Signatures in IPF Patients: A Multiplex Bead-Based Flow Cytometry Approach. Cells 10:

De Langhe, E., C. Aznar-Lopez, V. De Vooght, J.A. Vanoirbeek, F.P. Luyten, and R.J. Lories. 2014. Secreted frizzled related proteins inhibit fibrosis in vitro but appear redundant in vivo. Fibrogenesis Tissue Repair 7:14.

Distler, J.H.W., A.H. Gyorfi, M. Ramanujam, M.L. Whitfield, M. Konigshoff, and R. Lafyatis. 2019. Shared and distinct mechanisms of fibrosis. Nat Rev Rheumatol 15:705–730.

Elliot, S., P. Catanuto, S. Pereira-Simon, X. Xia, S. Shahzeidi, E. Roberts, J. Ludlow, S. Hamdan, S. Daunert, J. Parra, R. Stone, I. Pastar, M. Tomic-Canic, and M.K. Glassberg. 2022. Urine-derived exosomes from individuals with IPF carry pro-fibrotic cargo. Elife 11:

Elzi, D.J., M. Song, K. Hakala, S.T. Weintraub, and Y. Shiio. 2012. Wnt antagonist SFRP1 functions as a secreted mediator of senescence. Mol Cell Biol 32:4388–4399.

Englert, J.M., L.E. Hanford, N. Kaminski, J.M. Tobolewski, R.J. Tan, C.L. Fattman, L. Ramsgaard, T.J. Richards, I. Loutaev, P.P. Nawroth, M. Kasper, A. Bierhaus, and T.D. Oury. 2008. A role for the receptor for advanced glycation end products in idiopathic pulmonary fibrosis. Am J Pathol 172:583–591.

Fernandez, I.E., O.V. Amarie, K. Mutze, M. Konigshoff, A.O. Yildirim, and O. Eickelberg. 2016. Systematic phenotyping and correlation of biomarkers with lung function and histology in lung fibrosis. Am J Physiol Lung Cell Mol Physiol 310:L919–927.

Fernandez, I.E., and O. Eickelberg. 2012. New cellular and molecular mechanisms of lung injury and fibrosis in idiopathic pulmonary fibrosis. Lancet 380:680–688.

Guiot, J., M. Cambier, A. Boeckx, M. Henket, O. Nivelles, F. Gester, E. Louis, M. Malaise, F. Dequiedt, R. Louis, I. Struman, and M.S. Njock. 2020. Macrophage-derived exosomes attenuate fibrosis in airway epithelial cells through delivery of antifibrotic miR-142-3p. Thorax 75:870–881.

Hu, Y., J.P. Ng-Blichfeldt, C. Ota, C. Ciminieri, W. Ren, P.S. Hiemstra, J. Stolk, R. Gosens, and M. Konigshoff. 2020. Wnt/beta-catenin signaling is critical for regenerative potential of distal lung epithelial progenitor cells in homeostasis and emphysema. Stem Cells 38:1467–1478.

Jenkins, R.G., B.B. Moore, R.C. Chambers, O. Eickelberg, M. Konigshoff, M. Kolb, G.J. Laurent, C.B. Nanthakumar, M.A. Olman, A. Pardo, M. Selman, D. Sheppard, P.J. Sime, A.M. Tager, A.L. Tatler, V.J. Thannickal, E.S. White, A.T.S.A.o.R. Cell, and B. Molecular. 2017. An Official American Thoracic Society Workshop Report: Use of Animal Models for the Preclinical Assessment of Potential Therapies for Pulmonary Fibrosis. Am J Respir Cell Mol Biol 56:667–679.

Kadota, T., Y. Fujita, J. Araya, N. Watanabe, S. Fujimoto, H. Kawamoto, S. Minagawa, H. Hara, T. Ohtsuka, Y. Yamamoto, K. Kuwano, and T. Ochiya. 2021. Human bronchial epithelial cell-derived extracellular vesicle therapy for pulmonary fibrosis via inhibition of TGF-beta-WNT crosstalk. J Extracell Vesicles 10:e12124.

Kadota, T., Y. Yoshioka, Y. Fujita, J. Araya, S. Minagawa, H. Hara, A. Miyamoto, S. Suzuki, S. Fujimori, T. Kohno, T. Fujii, K. Kishi, K. Kuwano, and T. Ochiya. 2020. Extracellular Vesicles from Fibroblasts Induce Epithelial-Cell Senescence in Pulmonary Fibrosis. Am J Respir Cell Mol Biol 63:623–636.

Kang, J.H., M.Y. Jung, M. Choudhury, and E.B. Leof. 2020. Transforming growth factor beta induces fibroblasts to express and release the immunomodulatory protein PD-L1 into extracellular vesicles. FASEB J 34:2213–2226.

Kaur, G., K.P. Maremanda, M. Campos, H.S. Chand, F. Li, N. Hirani, M.A. Haseeb, D. Li, and I. Rahman. 2021. Distinct Exosomal miRNA Profiles from BALF and Lung Tissue of COPD and IPF Patients. Int J Mol Sci 22:

Kavanagh, E.L., S. Lindsay, M. Halasz, L.C. Gubbins, K. Weiner-Gorzel, M.H.Z. Guang, A. McGoldrick, E. Collins, M. Henry, A. Blanco-Fernandez, O.G. P, P. Fitzpatrick, M.J. Higgins, P. Dowling, and A. McCann. 2017. Protein and chemotherapy profiling of extracellular vesicles harvested from therapeutic induced senescent triple negative breast cancer cells. Oncogenesis 6:e388.

King, T.E., Jr., W.Z. Bradford, S. Castro-Bernardini, E.A. Fagan, I. Glaspole, M.K. Glassberg, E. Gorina, P.M. Hopkins, D. Kardatzke, L. Lancaster, D.J. Lederer, S.D. Nathan, C.A. Pereira, S.A. Sahn, R. Sussman, J.J. Swigris, P.W. Noble, and A.S. Group. 2014. A phase 3 trial of pirfenidone in patients with idiopathic pulmonary fibrosis. N Engl J Med 370:2083–2092.

Konigshoff, M., N. Balsara, E.M. Pfaff, M. Kramer, I. Chrobak, W. Seeger, and O. Eickelberg. 2008. Functional Wnt signaling is increased in idiopathic pulmonary fibrosis. PLoS One 3:e2142.

Konigshoff, M., M. Kramer, N. Balsara, J. Wilhelm, O.V. Amarie, A. Jahn, F. Rose, L. Fink, W. Seeger, L. Schaefer, A. Gunther, and O. Eickelberg. 2009. WNT1-inducible signaling protein-1 mediates pulmonary fibrosis in mice and is upregulated in humans with idiopathic pulmonary fibrosis. J Clin Invest 119:772–787.

Kurche, J.S., I.T. Stancil, J.E. Michalski, I.V. Yang, and D.A. Schwartz. 2022. Dysregulated Cell-Cell Communication Characterizes Pulmonary Fibrosis. Cells 11:

Lacy, S.H., C.F. Woeller, T.H. Thatcher, S.J. Pollock, E.M. Small, P.J. Sime, and R.P. Phipps. 2019. Activated Human Lung Fibroblasts Produce Extracellular Vesicles with Antifibrotic Prostaglandins. Am J Respir Cell Mol Biol 60:269–278.

Lehmann, M., Q. Hu, Y. Hu, K. Hafner, R. Costa, A. van den Berg, and M. Konigshoff. 2020. Chronic WNT/beta-catenin signaling induces cellular senescence in lung epithelial cells. Cell Signal 70:109588.

Makiguchi, T., M. Yamada, Y. Yoshioka, H. Sugiura, A. Koarai, S. Chiba, N. Fujino, Y. Tojo, C. Ota, H. Kubo, S. Kobayashi, M. Yanai, S. Shimura, T. Ochiya, and M. Ichinose. 2016. Serum extracellular vesicular miR-21-5p is a predictor of the prognosis in idiopathic pulmonary fibrosis. Respir Res 17:110.

Mao, X., S.K. Tey, C.L.S. Yeung, E.M.L. Kwong, Y.M.E. Fung, C.Y.S. Chung, L.Y. Mak, D.K.H. Wong, M.F. Yuen, J.C.M. Ho, H. Pang, M.P. Wong, C.O. Leung, T.K.W. Lee, V. Ma, W.C. Cho, P. Cao, X. Xu, Y. Gao, and J.W.P. Yam. 2020. Nidogen 1-Enriched Extracellular Vesicles Facilitate Extrahepatic Metastasis of Liver Cancer by Activating Pulmonary Fibroblasts to Secrete Tumor Necrosis Factor Receptor 1. Adv Sci (Weinh) 7:2002157.

Martin-Medina, A., M. Lehmann, O. Burgy, S. Hermann, H.A. Baarsma, D.E. Wagner, M.M. De Santis, F. Ciolek, T.P. Hofer, M. Frankenberger, M. Aichler, M. Lindner, W. Gesierich, A. Guenther, A. Walch, C. Coughlan, P. Wolters, J.S. Lee, J. Behr, and M. Konigshoff. 2018. Increased Extracellular Vesicles Mediate WNT5A Signaling in Idiopathic Pulmonary Fibrosis. Am J Respir Crit Care Med 198:1527–1538.

Martinez, F.J., H.R. Collard, A. Pardo, G. Raghu, L. Richeldi, M. Selman, J.J. Swigris, H. Taniguchi, and A.U. Wells. 2017. Idiopathic pulmonary fibrosis. Nat Rev Dis Primers 3:17074.

Matsuyama, M., A. Nomori, K. Nakakuni, A. Shimono, and M. Fukushima. 2014. Secreted Frizzled-related protein 1 (Sfrp1) regulates the progression of renal fibrosis in a mouse model of obstructive nephropathy. J Biol Chem 289:31526–31533.

Mayr, C.H., A. Sengupta, M. Ansari, J.C. Pestoni, P. Ogar, I. Angelidis, A. Liontos, A. Rodriguez-Castillo, N.J. Lang, M. Strunz, S. Asgharpour, D. Porras-Gonzalez, M. Gerckens, B. Oehrle, V. Viteri-Alvarez, I.E. Fernandez, M. Tallquist, M. Irmler, J. Beckers, O. Eickelberg, G. Mircea Stoleriu, J. Behr, N. Kneidinger, A.Ö. Yildirim, K. Ahlbrecht, R.E. Morty, C. Samakovlis, F.J. Theis, G. Burgstaller, and H.B. Schiller. 2022. Autocrine Sfrp1 inhibits lung fibroblast invasion during transition to injury induced myofibroblasts. BioRxiv

Mayr, C.H., L.M. Simon, G. Leuschner, M. Ansari, J. Schniering, P.E. Geyer, I. Angelidis, M. Strunz, P. Singh, N. Kneidinger, F. Reichenberger, E. Silbernagel, S. Bohm, H. Adler, M. Lindner, B. Maurer, A. Hilgendorff, A. Prasse, J. Behr, M. Mann, O. Eickelberg, F.J. Theis, and H. B. Schiller. 2021. Integrative analysis of cell state changes in lung fibrosis with peripheral protein biomarkers. EMBO Mol Med 13:e12871.

Mummler, C., O. Burgy, S. Hermann, K. Mutze, A. Gunther, and M. Konigshoff. 2018. Cellspecific expression of runt-related transcription factor 2 contributes to pulmonary fibrosis. FASEB J 32:703–716.

Mutze, K., S. Vierkotten, J. Milosevic, O. Eickelberg, and M. Konigshoff. 2015. Enolase 1 (ENO1) and protein disulfide-isomerase associated 3 (PDIA3) regulate Wnt/beta-catenin-driven trans-differentiation of murine alveolar epithelial cells. Dis Model Mech 8:877–890.

Ng-Blichfeldt, J.P., T. de Jong, R.K. Kortekaas, X. Wu, M. Lindner, V. Guryev, P.S. Hiemstra, J. Stolk, M. Konigshoff, and R. Gosens. 2019. TGF-beta activation impairs fibroblast ability to support adult lung epithelial progenitor cell organoid formation. Am J Physiol Lung Cell Mol Physiol 317:L14–L28.

Njock, M.S., J. Guiot, M.A. Henket, O. Nivelles, M. Thiry, F. Dequiedt, J.L. Corhay, R.E. Louis, and I. Struman. 2019. Sputum exosomes: promising biomarkers for idiopathic pulmonary fibrosis. Thorax 74:309–312.

Parimon, T., C. Yao, D.M. Habiel, L. Ge, S.A. Bora, R. Brauer, C.M. Evans, T. Xie, F. Alonso-Valenteen, L.K. Medina-Kauwe, D. Jiang, P.W. Noble, C.M. Hogaboam, N. Deng, O. Burgy, T.J. Antes, M. Konigshoff, B.R. Stripp, S.A. Gharib, and P. Chen. 2019. Syndecan-1 promotes lung fibrosis by regulating epithelial reprogramming through extracellular vesicles. JCI Insight 5:

Patterson, S.A., G. Deep, and T.E. Brinkley. 2018. Detection of the receptor for advanced glycation endproducts in neuronally-derived exosomes in plasma. Biochem Biophys Res Commun 500:892–896.

Richeldi, L., V. Cottin, R.M. du Bois, M. Selman, T. Kimura, Z. Bailes, R. Schlenker-Herceg, S. Stowasser, and K.K. Brown. 2016. Nintedanib in patients with idiopathic pulmonary fibrosis: Combined evidence from the TOMORROW and INPULSIS((R)) trials. Respir Med 113:74–79.

Santos-Alvarez, J.C., J.M. Velazquez-Enriquez, R. Garcia-Carrillo, C. Rodriguez-Beas, A.A. Ramirez-Hernandez, E. Reyes-Jimenez, K. Gonzalez-Garcia, A. Lopez-Martinez, L. Perez-Campos Mayoral, S.R. Aguilar-Ruiz, M.L.A. Romero-Tlalolini, H. Torres-Aguilar, L. Castro-Sanchez, J. Arellanes-Robledo, V.R. Vasquez-Garzon, and R. Baltierrez-Hoyos. 2022. miRNAs Contained in Extracellular Vesicles Cargo Contribute to the Progression of Idiopathic Pulmonary Fibrosis: An In Vitro Aproach. Cells 11:

Satoh, W., T. Gotoh, Y. Tsunematsu, S. Aizawa, and A. Shimono. 2006. Sfrp1 and Sfrp2 regulate anteroposterior axis elongation and somite segmentation during mouse embryogenesis. Development 133:989–999.

Schiller, H.B., I.E. Fernandez, G. Burgstaller, C. Schaab, R.A. Scheltema, T. Schwarzmayr, T. M. Strom, O. Eickelberg, and M. Mann. 2015. Time- and compartment-resolved proteome profiling of the extracellular niche in lung injury and repair. Mol Syst Biol 11:819.

Shaba, E., C. Landi, A. Carleo, L. Vantaggiato, E. Paccagnini, M. Gentile, L. Bianchi, P. Lupetti, E. Bargagli, A. Prasse, and L. Bini. 2021. Proteome Characterization of BALF Extracellular Vesicles in Idiopathic Pulmonary Fibrosis: Unveiling Undercover Molecular Pathways. Int J Mol Sci 22:

Soekmadji, C., B. Li, Y. Huang, H. Wang, T. An, C. Liu, W. Pan, J. Chen, L. Cheung, J.M. Falcon-Perez, Y.S. Gho, H.B. Holthofer, M.T.N. Le, A. Marcilla, L. O’Driscoll, F. Shekari, T.L. Shen, A.C. Torrecilhas, X. Yan, F. Yang, H. Yin, Y. Xiao, Z. Zhao, X. Zou, Q. Wang, and L. Zheng. 2020. The future of Extracellular Vesicles as Theranostics - an ISEV meeting report. J Extracell Vesicles 9:1809766.

Strunz, M., L.M. Simon, M. Ansari, J.J. Kathiriya, I. Angelidis, C.H. Mayr, G. Tsidiridis, M. Lange, L.F. Mattner, M. Yee, P. Ogar, A. Sengupta, I. Kukhtevich, R. Schneider, Z. Zhao, C. Voss, T. Stoeger, J.H.L. Neumann, A. Hilgendorff, J. Behr, M. O’Reilly, M. Lehmann, G. Burgstaller, M. Konigshoff, H.A. Chapman, F.J. Theis, and H.B. Schiller. 2020. Alveolar regeneration through a Krt8+ transitional stem cell state that persists in human lung fibrosis. Nat Commun 11:3559.

Thery, C., K.W. Witwer, E. Aikawa, M.J. Alcaraz, J.D. Anderson, R. Andriantsitohaina, A. Antoniou, T. Arab, F. Archer, G.K. Atkin-Smith, D.C. Ayre, J.M. Bach, D. Bachurski, H. Baharvand, L. Balaj, S. Baldacchino, N.N. Bauer, A.A. Baxter, M. Bebawy, C. Beckham, A. Bedina Zavec, A. Benmoussa, A.C. Berardi, P. Bergese, E. Bielska, C. Blenkiron, S. Bobis-Wozowicz, E. Boilard, W. Boireau, A. Bongiovanni, F.E. Borras, S. Bosch, C.M. Boulanger, X. Breakefield, A.M. Breglio, M.A. Brennan, D.R. Brigstock, A. Brisson, M.L. Broekman, J.F. Bromberg, P. Bryl-Gorecka, S. Buch, A.H. Buck, D. Burger, S. Busatto, D. Buschmann, B. Bussolati, E.I. Buzas, J.B. Byrd, G. Camussi, D.R. Carter, S. Caruso, L.W. Chamley, Y.T. Chang, C. Chen, S. Chen, L. Cheng, A.R. Chin, A. Clayton, S.P. Clerici, A. Cocks, E. Cocucci, R.J. Coffey, A. Cordeiro-da-Silva, Y. Couch, F.A. Coumans, B. Coyle, R. Crescitelli, M.F. Criado, C. D’Souza-Schorey, S. Das, A. Datta Chaudhuri, P. de Candia, E.F. De Santana, O. De Wever, H.A. Del Portillo, T. Demaret, S. Deville, A. Devitt, B. Dhondt, D. Di Vizio, L.C. Dieterich, V. Dolo, A.P. Dominguez Rubio, M. Dominici, M.R. Dourado, T.A. Driedonks, F.V. Duarte, H.M. Duncan, R.M. Eichenberger, K. Ekstrom, S. El Andaloussi, C. Elie-Caille, U. Erdbrugger, J.M. Falcon-Perez, F. Fatima, J.E. Fish, M. Flores-Bellver, A. Forsonits, A. Frelet-Barrand, F. Fricke, G. Fuhrmann, S. Gabrielsson, A. Gamez-Valero, C. Gardiner, K. Gartner, R. Gaudin, Y.S. Gho, B. Giebel, C. Gilbert, M. Gimona, I. Giusti, D.C. Goberdhan, A. Gorgens, S. M. Gorski, D.W. Greening, J.C. Gross, A. Gualerzi, G.N. Gupta, D. Gustafson, A. Handberg, R.A. Haraszti, P. Harrison, H. Hegyesi, A. Hendrix, A.F. Hill, F.H. Hochberg, K.F. Hoffmann, B. Holder, H. Holthofer, B. Hosseinkhani, G. Hu, Y. Huang, V. Huber, S. Hunt, A.G. Ibrahim, T. Ikezu, J.M. Inal, M. Isin, A. Ivanova, H.K. Jackson, S. Jacobsen, S.M. Jay, M. Jayachandran, G. Jenster, L. Jiang, S.M. Johnson, J.C. Jones, A. Jong, T. Jovanovic-Talisman, S. Jung, R. Kalluri, S.I. Kano, S. Kaur, Y. Kawamura, E.T. Keller, D. Khamari, E. Khomyakova, A. Khvorova, P. Kierulf, K.P. Kim, T. Kislinger, M. Klingeborn, D.J. Klinke, 2nd, M. Kornek, M.M. Kosanovic, A.F. Kovacs, E.M. Kramer-Albers, S. Krasemann, M. Krause, I.V. Kurochkin, G.D. Kusuma, S. Kuypers, S. Laitinen, S.M. Langevin, L.R. Languino, J. Lannigan, C. Lasser, L.C. Laurent, G. Lavieu, E. Lazaro-Ibanez, S. Le Lay, M.S. Lee, Y.X.F. Lee, D.S. Lemos, M. Lenassi, A. Leszczynska, I.T. Li, K. Liao, S.F. Libregts, E. Ligeti, R. Lim, S.K. Lim, A. Line, K. Linnemannstons, A. Llorente, C.A. Lombard, M.J. Lorenowicz, A.M. Lorincz, J. Lotvall, J. Lovett, M.C. Lowry, X. Loyer, Q. Lu, B. Lukomska, T.R. Lunavat, S.L. Maas, H. Malhi, A. Marcilla, J. Mariani, J. Mariscal, E.S. Martens-Uzunova, L. Martin-Jaular, M.C. Martinez, V.R. Martins, M. Mathieu, S. Mathivanan, M. Maugeri, L.K. McGinnis, M.J. McVey, D. G. Meckes, Jr., K.L. Meehan, I. Mertens, V.R. Minciacchi, A. Moller, M. Moller Jorgensen, A. Morales-Kastresana, J. Morhayim, F. Mullier, M. Muraca, L. Musante, V. Mussack, D.C. Muth, K.H. Myburgh, T. Najrana, M. Nawaz, I. Nazarenko, P. Nejsum, C. Neri, T. Neri, R. Nieuwland, L. Nimrichter, J.P. Nolan, E.N. Nolte-’t Hoen, N. Noren Hooten, L. O’Driscoll, T. O’Grady, A. O’Loghlen, T. Ochiya, M. Olivier, A. Ortiz, L.A. Ortiz, X. Osteikoetxea, O. Ostergaard, M. Ostrowski, J. Park, D.M. Pegtel, H. Peinado, F. Perut, M.W. Pfaffl, D.G. Phinney, B.C. Pieters, R.C. Pink, D.S. Pisetsky, E. Pogge von Strandmann, I. Polakovicova, I.K. Poon, B.H. Powell, I. Prada, L. Pulliam, P. Quesenberry, A. Radeghieri, R.L. Raffai, S. Raimondo, J. Rak, M.I. Ramirez, G. Raposo, M.S. Rayyan, N. Regev-Rudzki, F.L. Ricklefs, P.D. Robbins, D.D. Roberts, S.C. Rodrigues, E. Rohde, S. Rome, K.M. Rouschop, A. Rughetti, A.E. Russell, P. Saa, S. Sahoo, E. Salas-Huenuleo, C. Sanchez, J.A. Saugstad, M.J. Saul, R. M. Schiffelers, R. Schneider, T.H. Schoyen, A. Scott, E. Shahaj, S. Sharma, O. Shatnyeva, F. Shekari, G.V. Shelke, A.K. Shetty, K. Shiba, P.R. Siljander, A.M. Silva, A. Skowronek, O.L. Snyder, 2nd, R.P. Soares, B.W. Sodar, C. Soekmadji, J. Sotillo, P.D. Stahl, W. Stoorvogel, S. L. Stott, E.F. Strasser, S. Swift, H. Tahara, M. Tewari, K. Timms, S. Tiwari, R. Tixeira, M. Tkach, W.S. Toh, R. Tomasini, A.C. Torrecilhas, J.P. Tosar, V. Toxavidis, L. Urbanelli, P. Vader, B.W. van Balkom, S.G. van der Grein, J. Van Deun, M.J. van Herwijnen, K. Van Keuren-Jensen, G. van Niel, M.E. van Royen, A.J. van Wijnen, M.H. Vasconcelos, I.J. Vechetti, Jr., T.D. Veit, L.J. Vella, E. Velot, F.J. Verweij, B. Vestad, J.L. Vinas, T. Visnovitz, K.V. Vukman, J. Wahlgren, D.C. Watson, M.H. Wauben, A. Weaver, J.P. Webber, V. Weber, A.M. Wehman, D.J. Weiss, J.A. Welsh, S. Wendt, A.M. Wheelock, Z. Wiener, L. Witte, J. Wolfram, A. Xagorari, P. Xander, J. Xu, X. Yan, M. Yanez-Mo, H. Yin, Y. Yuana, V. Zappulli, J. Zarubova, V. Zekas, J.Y. Zhang, Z. Zhao, L. Zheng, A.R. Zheutlin, A.M. Zickler, P. Zimmermann, A.M. Zivkovic, D. Zocco, and E.K. Zuba-Surma. 2018. Minimal information for studies of extracellular vesicles 2018 (MISEV2018): a position statement of the International Society for Extracellular Vesicles and update of the MISEV2014 guidelines. J Extracell Vesicles 7:1535750.

Tyanova, S., T. Temu, P. Sinitcyn, A. Carlson, M.Y. Hein, T. Geiger, M. Mann, and J. Cox. 2016. The Perseus computational platform for comprehensive analysis of (prote)omics data. Nat Methods 13:731–740.

Tzouvelekis, A., P. Ntolios, A. Karameris, G. Vilaras, P. Boglou, A. Koulelidis, K. Archontogeorgis, K. Kaltsas, G. Zacharis, E. Sarikloglou, P. Steiropoulos, D. Mikroulis, A. Koutsopoulos, M. Froudarakis, and D. Bouros. 2013. Increased expression of epidermal growth factor receptor (EGF-R) in patients with different forms of lung fibrosis. Biomed Res Int 2013:654354.

Uhl, F.E., S. Vierkotten, D.E. Wagner, G. Burgstaller, R. Costa, I. Koch, M. Lindner, S. Meiners, O. Eickelberg, and M. Konigshoff. 2015. Preclinical validation and imaging of Wnt-induced repair in human 3D lung tissue cultures. Eur Respir J 46:1150–1166.

van Niel, G., D.R.F. Carter, A. Clayton, D.W. Lambert, G. Raposo, and P. Vader. 2022. Challenges and directions in studying cell-cell communication by extracellular vesicles. Nat Rev Mol Cell Biol 23:369–382.

van Niel, G., G. D’Angelo, and G. Raposo. 2018. Shedding light on the cell biology of extracellular vesicles. Nat Rev Mol Cell Biol 19:213–228.

Vuga, L.J., A. Ben-Yehudah, E. Kovkarova-Naumovski, T. Oriss, K.F. Gibson, C. Feghali-Bostwick, and N. Kaminski. 2009. WNT5A is a regulator of fibroblast proliferation and resistance to apoptosis. Am J Respir Cell Mol Biol 41:583–589.

Wolf, F.A., P. Angerer, and F.J. Theis. 2018. SCANPY: large-scale single-cell gene expression data analysis. Genome Biol 19:15.

Wynn, T.A. 2007. Common and unique mechanisms regulate fibrosis in various fibroproliferative diseases. J Clin Invest 117:524–529.

Xie, T., V. Kulur, N. Liu, N. Deng, Y. Wang, S.C. Rowan, C. Yao, G. Huang, X. Liu, F. Taghavifar, J. Liang, C. Hogaboam, B. Stripp, P. Chen, D. Jiang, and P.W. Noble. 2021. Mesenchymal growth hormone receptor deficiency leads to failure of alveolar progenitor cell function and severe pulmonary fibrosis. Sci Adv 7:

Zhang, H., T. Deng, R. Liu, M. Bai, L. Zhou, X. Wang, S. Li, X. Wang, H. Yang, J. Li, T. Ning, D. Huang, H. Li, L. Zhang, G. Ying, and Y. Ba. 2017. Exosome-delivered EGFR regulates liver microenvironment to promote gastric cancer liver metastasis. Nat Commun 8:15016.

Zhou, B., K. Xu, X. Zheng, T. Chen, J. Wang, Y. Song, Y. Shao, and S. Zheng. 2020. Application of exosomes as liquid biopsy in clinical diagnosis. Signal Transduct Target Ther 5:144.

Zomer, A., C. Maynard, F.J. Verweij, A. Kamermans, R. Schafer, E. Beerling, R.M. Schiffelers, E. de Wit, J. Berenguer, S.I.J. Ellenbroek, T. Wurdinger, D.M. Pegtel, and J. van Rheenen. 2015. In Vivo imaging reveals extracellular vesicle-mediated phenocopying of metastatic behavior. Cell 161:1046–1057.

